# Exploring The Formation And Permeability Of Plasmodesmata In The Liverwort, *Marchantia polymorpha*

**DOI:** 10.1101/2024.12.16.628828

**Authors:** Chia-Yun Hsu, Chia-Hsuan Hsu, Hui-Yu Chang, Kuan-Ju Lu

**Affiliations:** Graduate Institute of Biotechnology, National Chung Hsing University. 145 Xingda Rd., South Dist., Taichung City 40227, Taiwan, R.O.C; Advanced Plant and Food Crop Biotechnology Center, National Chung Hsing University, Taichung, 402, Taiwan

**Keywords:** callose, *Marchantia polymorpha*, particle bombardment, photo-conversion, plasmodesmata

## Abstract

Plasmodesmata are cell-wall-embedded channels that evolved in the common ancestor of land plants to increase cell-to-cell communication. Whether all the fundamental properties of plasmodesmata emerged and were inherited in all land plants at the same time is unknown. Here we show that the bryophyte *Marchantia polymorpha* (a non-vascular plant) forms mostly simple plasmodesmata in early-developing gemmae. The complexity of plasmodesmata increases during gemma maturation, and complex plasmodesmata with enlarged cavities are majorly observed in thalli. In contrast to vascular plants, whose simple plasmodesmata can transport monomeric fluorescent proteins, plasmodesmata in *Marchantia polymorpha* limited their permeability before the juvenile-to-adult transition. In support, callose, a known polysaccharide regulating plasmodesmata permeability in vascular plants, accumulated in most of the *Marchantia polymorpha* tissues examined. Furthermore, we found that in the apical meristematic region, plasmodesmata allowed the transport of monomeric fluorescent proteins, and this relaxation might correlate with the lower accumulation of callose. Taken together, our study suggests that certain plasmodesmata properties, such as complexity progression and callose accumulation, may have evolved before the divergence between vascular and non-vascular plants.

## Introduction

Land plants transitioned from water to land approximately 450 million years ago. Following terrestrialization, plants underwent numerous innovations, including the development of stomata, surface protections on the epidermis, and rhizoids, enabling them to adapt to drastic environmental changes (Bowman et al. 2017; Cheng et al. 2019; Li et al. 2020; Zhang et al. 2020). Among these innovations, the emergence of plasmodesmata marks a crucial event in the evolution of higher-order multicellular bodies. In the past 50 years, it has been well demonstrated that plasmodesmata are not only responsible for the transport of molecules but also involved in the determination of cell fates, development of organs, and response to biotic and abiotic stresses (Bayer and Benitez-Alfonso 2024; Liu et al. 2024; Lu et al. 2018; Reagan and Burch-Smith 2020). It was proposed that plasmodesmata-like structures evolved independently at least six times during the Plantae evolutionary process, and their emergence is highly correlated with the increase of the organism’s complexity (Brunkard and Zambryski 2016; Raven 2007). With the extensive genome decoding of many algae and land plants, the evolution of plasmodesmata might have some room for discussion. However, the structural differences between algal and land plant plasmodesmata still support the idea that plasmodesmata in land plants originated as a single event in the most common ancestor of land plants, predating the divergence of bryophytes (non-vascular plants) and Tracheophyta (vascular plants) (Brunkard and Zambryski 2016; Raven 2007). Plasmodesmata have been inherited by all land plants, underscoring the indispensable role of plasmodesmata in terrestrial lifestyles.

Plasmodesmata are membrane-lined nanochannels that connect most of the cells in plants (Bayer and Benitez-Alfonso 2024; Brunkard and Zambryski 2016; Li et al. 2021; Sager and Lee 2018). Plants utilize plasmodesmata to transport not only small molecules such as ions and sugars but also larger molecules like RNAs and proteins, facilitating coordinated physiological responses between cells (Gallagher et al. 2014; Liu and Chen 2018; Long et al. 2015; Lu et al. 2018). The size exclusion limit (SEL), representing the maximum size of molecules passively transported through plasmodesmata, was a focal point in early plasmodesmata research. Micro-injection studies, introducing fluorescent-labeled dextrans of varying molecular weights to cells, revealed that the SEL of PD is less than 10 KDa (Wolf et al. 1989). Subsequent research, involving the expression of fluorescent proteins through particle bombardment and tissue-specific promoters, demonstrated that the SEL in sink tissues can reach approximately 50-60 KDa. However, the SEL decreases to below 10 KDa after the transition from sink to source tissues (Oparka et al. 1999). It is widely accepted that the SEL varies among different tissues and is tightly regulated during development (Li et al. 2021).

One of the known plasmodesmata permeability regulations involves the dynamic accumulation of callose at the neck regions of plasmodesmata (Amsbury et al. 2017; De Storme and Geelen 2014; Levy et al. 2007; Sager and Lee 2018; Vaten et al. 2011; Wu et al. 2018). When plants face environmental or biological challenges, callose accumulation increases to physically block the plasmodesmata’s aperture (Amsbury et al. 2017; De Storme and Geelen 2014; Sager and Lee 2018; Wu et al. 2018). Callose synthases (CalSs) are responsible for the synthesis of callose, while β-1,3-glucanases (BGs) are responsible for its degradation (Amsbury et al. 2017; De Storme and Geelen 2014; Sager and Lee 2018; Wu et al. 2018). During development, plants must finely tune the SEL of plasmodesmata to prevent mobile signals from activating developmental processes under unfavorable conditions. In perennial trees, dormancy is a key process for surviving harsh winters. The accumulation of callose surrounding the shoot apical meristem (SAM) during winter inhibits the growth of buds (Rinne et al. 2001). When optimal conditions arrive during spring, plants degrade the callose, allowing awakening signals to be transported into the buds (Rinne et al. 2011). Additionally, recent studies indicate that tight regulation of plasmodesmata permeability is essential for phototropism of the hypocotyl (Han et al. 2014), the patterning of lateral root formation (Benitez-Alfonso et al. 2013), correct vein patterning of leaves (Linh and Scarpella 2022), and proper auxin distribution in root (Mellor et al. 2020) demonstrating the importance of regulating plasmodesmata permeability during development.

As plasmodesmata of land plants evolved before the divergence of bryophytes and vascular plants (Brunkard and Zambryski 2019; Raven 2007), it seems possible that a core mechanism that controls their permeability evolved prior to the divergence. Intercellular communication has been reported in one bryophyte, the moss *Physcomitrium patens* (previously *Physcomitrella patens*) (Kitagawa and Fujita 2015). The spread of fluorescence into neighboring cells was observed within 30 minutes after inducing a photo-inducible fluorescent protein illumination in one of the filamentous protonema cells (Kitagawa and Fujita 2015). The transport of the fluorescent protein could be interrupted by applying the stress-induced plant hormone abscisic acid (ABA). Mutants that abolish the ABA synthesis also exhibit increased plasmodesmata permeability (Kitagawa et al. 2019). The aperture of plasmodesmata is narrower in ABA-treated wild-type plants (WT) compared to mock-treated WT and ABA-treated *aba1* mutants, suggesting that moss may respond to stress through the ABA signaling pathway to reduce the intercellular communication (Kitagawa et al. 2019). However, how bryophytes coordinate development and PD permeability remains unclear.

The complete genome information reveals that most key protein families are present and with less redundancy when compared to most vascular model plants in one of the model bryophytes, *Marchantia polymorpha* (hereafter referred to as *M. polymorpha*). In comparison to another bryophyte, *Physcomitrium patens*, *M. polymorpha* did not undergo whole-genome duplication. Therefore, many molecular pathways and the composition of protein complexes are easier to explore. For functional analyses, M. polumorpha will provde great advatages (Cesarino et al. 2020). Additionally, compared to the predominantly two-dimensional growth of *P. patens*, the simple three-dimensional structure of *M. polymorpha* may allow us to investigate how lateral and horizontal intercellular communication is coordinated during evolution. These properties encourages us to explore the divergency of plasmodesmata properties in *M. polymorpha* and may establish the system as an alternative model system for bryophytic plasmodesmata research.

Our objective is to understand how *M. polymorpha* regulates plasmodesmata during development. We found that *M. polymorpha* majorly forms simple plasmodesmata in early-developing gemmae, progressing to more complex plasmodesmata during the gemmae maturation and turning into more complex plasmodesmata with large cavities in thalli. In correlation with the limited intercellular mobility of a photo-convertible mEos3.2 fluorescent protein, dormant gemmae accumulated a significant amount of callose surrounding most cells. This permeability limitation persisted even after the germination of gemmae. This observation also holds for thalli, aligning well with the maintained callose accumulation. However, we noted a broader traffic range of superfolder yellow fluorescent protein 2 (SYFP2) when bombarding thalli, compared to gemmalings. This suggests that *M. polymorpha* may regulate the increase in SEL when necessray. Finally, through tissue-specific expression, we demonstrated that, in accord with low callose accumulation, plasmodesmata are more permeable around the apical notch. Our study unveils the fundamental properties of plasmodesmata and suggests that callose accumulation might regulate PD permeability in *M. polymorpha*.

## Material and method

### Electron microscopy

Gemmae and gemma cups were collected from 21-day-old thalli, and samples were fixed in 2.5% glutaraldehyde and 3.2% paraformaldehyde in 0.1 M sodium phosphate buffer, pH 7.0. For post-fixation treatment, samples were transferred to 1% OsO₄ in the same 0.1 M sodium phosphate buffer, pH 7.0, followed by dehydration with an ethanol series (from 15% to 100%) and 100% propylene oxide. Spurr resins were used for infiltration and embedding. The Leica EM UC7 ultramicrotome was used for the ultra-thin section (70–90 nm). The sections were stained with 5% uranyl acetate in 50% methanol and 0.4% lead citrate in 0.1 N sodium hydroxide. One thin slide of sections was stained with Toluidine blue and observed under a stereo microscope to record the outline of the cells. All samples were observed using a FEI G2 Tecnai Spirit Twin transmission electron microscope at 80 kV.

### Constructions

*p35S::mEos3.2* construct was cloned by SLiCE reaction and Gateway^®^ system (Invitrogen). The MpGWB system (Ishizaki et al. 2015) was used as the final destination vector for plant transformation. The SLiCE buffer and SLiCE extract were used as previously reported with some adjustments (Zhang et al., 2012). The SLiCE extract was produced from the TOP10 *E. coli* strain. We modified the pMpGWB102 to fit with the SLiCE method. A 30nt ligation-independent cloning (LIC) linker fragment was synthesized based on a previous publication (Table S1; De Rybel et al. 2011) and cloned into the pENTR-D-TOPO^®^ vector following the supplier’s instructions (Invitrogen) to generate a pENTR-LIC plasmid. Following the supplier’s recommendation, the LIC fragment was sub-cloned into pMpGWB102 using the LR Clonase™ II Plus enzyme (Invitrogen). mEos3.2 was PCR-amplified from the mEos3.2-N1 plasmid using primers containing a 15nt linker fragment homologous to the LIC site (Table S1) and the pMpGWB-LIC vectors were linearized by HpaI digestion. For the SLiCE reaction, a 50-100 ng linear vector was mixed with insert DNA in a 3:1 (insert: vector) molar ratio. The SLiCE reaction was incubated at 37°C for 1h and chemically transformed into ECOS™101 DH5α competent cells (Yeastern Biotech, Taiwan) following the manufacturer’s instructions. mEos3.2-N1 was a gift from Michael Davidson & Tao Xu (Addgene plasmid # 54525; http://n2t.net/addgene:54525; RRID: Addgene_54525). pMpGWB102 was a gift from Takayuki Kohchi (Addgene plasmid # 68556; http://n2t.net/addgene:68556; RRID: Addgene_68556)

*MpLAXRpro::tdT-nls* and pENTR-*MpLAXRpro* were kindly provided by Ryuichi Nishihama (Ishida et al. 2022). *MpLAXRpro::mCitrine* construct was generated by transferring the *MpLARXpro* fragment from pENTR into the pMpGWB107 vector with the LR Clonase™ II Plus enzyme (Invitrogen) as the supplier’s recommendation. pMpGWB107 was a gift from Takayuki Kohchi (Addgene plasmid # 68560; http://n2t.net/addgene:68560; RRID: Addgene_68560)

The pMON *35S::LIC-SYFP2* plasmid used in the particle bombardments was kindly provided by Dolf Weijers (Yoshida et al. 2019).

### Plant growth condition

Gemmae from male (Takaragaike-1; Tak-1) *M. polymorpha* were cultured on 1/2 B5 medium [1/2 Gamborg’s B5 medium (Duchefa), 0.5 g/L MES, pH 5.7 and 1% agar], and placed in growth chamber (F-740, HiPoint), at 22°C under a 16h/8h day/night cycles, with 60 μEm^−2^s^−1^ LED white light.

*N. benthamiana* was grown in 10 cm pots in a mix of equal volumes of potting mixture and vermiculite at 25°C under long-day conditions (16 h/8 h light/dark cycles).

### Plant transformation

Transformation of *M. polymorpha* is performed according to the published method (Kubota et al. 2013). In brief, remove the apical meristem area of 14-day-old thalli, cut the thallus into small pieces, and incubate on 1/2 B5 medium with sucrose (1/2 Gamborg’s B5 medium, 0.5 g/L MES, pH 5.7, 1% agar, and 1% sucrose) for 3 days. The tissues were then cocultured with OD_600_ = 1.0 agrobacterium in 0M51C medium supplied with 2% sucrose and 100 μM acetosyringone (3,5-dimethoxy-4-hydroxyacetophenone) under a 16h/8h day/night cycle with agitation at 100 rpm at 22°C for another 3 days. The cultured tissues were washed with ddH_2_O several times and transferred onto 1/2 B5 medium containing antibiotics for selection.

### Confocal microscopy

Photoconversion confocal images were taken by the Leica Stellaris 8 confocal system in photon counting mode with the white light laser. mEos3.2 before photo-conversion images were taken with 50% 488 nm excitation and 500-550 nm detection and 75% 561 nm excitation and 600-650 nm detection for mEos3.2 after photo-conversion. All images were taken in photon counting mode with the gating set to collect photons from 1 nanosecond (ns) to 10 ns. Aniline blue staining and SYFP2 mobility images were taken by the Olympus FV3000 confocal system. The 405 nm diode laser (100 mW output) was set to 1%, and the detection wavelength was set between 450-500 nm for anilie blue. 5% 514 nm excitation and 550-600 nm detection was used for SYFP2. The bright field image of the mature thallus was generated by the imageJ stack focuser plug-in (https://imagej.net/ij/plugins/stack-focuser.html)

### Aniline blue staining

For gemmae staining, 10 μl of 0.1 mg/ml aniline blue fluorochrome (Biosupplies) was applied onto a slide. A few gemmae were transferred onto the droplet, covered by a cover slide, incubated for 10 minutes, and then imaged under the FV3000 confocal microscope.

For thalli, a concave slide was used to hold the 14-day-old thallus. 10 μl of 0.1 mg/ml aniline blue fluorochrome was directly applied onto the surface of the thallus and incubated for 10 minutes. The specimen was directly imaged under the FV3000 confocal microscope.

### Photoconversion

Photoconversion was performed using the Leica Stellaris confocal system. 1% of the 405 nm diode laser was selected for inducing conversion. For whole gemmae induction, the free running mode with a 400 Hz scanning speed was used to scan the entire focal plane, and the setting for mEos3.2 emission was as described previously. For single-cell induction, the region of interest was selected using the zoom-in function. The induction was performed by scanning the region with the 405 nm diode laser set to 1%, under 512 x 512 pixels, and a 400 Hz scanning speed for 30 seconds. Images before and after induction were taken with the settings described previously.

### Particle bombardment

The preparation of the gold particle was done according to the instruction manual for the Biolistic PDS-1000/He (Bio-Rad) device. 1 μg of plasmid DNA was mixed with 60 mg gold particles (1 μm in diameter) and 50 μl 2.5 M CaCl_2_, and vortexed briefly; 20 μl of 0.1 M spermidine was then added and vortexed. Gold particles were precipitated by centrifugation. The pellet was washed and resuspended in 60 μl 100% EtOH. 10 μl of the final gold particles were used for one transformation.

For the *N. bethamiana* experiment, 7 leaf discs were taken using a paper puncher and placed on a 3 cm 1/2 B5 medium agar plate. The samples were bombarded with gold particles at 900 psi from a 10 cm distance in the chamber with a 25 Hg vacuum. For *M. polymorpha* experiments, gemmae were picked by forceps from the gemma cup and placed concentrically on a 3 cm 1/2 B5 medium agar plate to form a 1 cm circle. For the 14-day-old thallus experiment, 3 thalli were picked by forceps and placed in the center of a 3 cm 1/2 B5 medium agar plate. The samples were bombarded with gold particles at 1100 psi from a 10 cm distance in the chamber with a 25 Hg vacuum. The bombarded samples were kept in the dark and left in a 21 °C growth chamber for 48 hours before observation.

## Results

### Dormant gemmae harbor primarily simple and few complex plasmodesmata

Plasmodesmata are categorized into two types in vascular plants based on their formation processes: primary and secondary plasmodesmata. Primary plasmodesmata are established during cytokinesis, and the majority exhibit a simple structure with one opening on each side of a cell wall. Conversely, secondary plasmodesmata are formed after cell division through an ambiguous mechanism (Burch-Smith et al. 2011). It has been suggested that secondary plasmodesmata predominantly arise near primary plasmodesmata, often developing into relatively complex structures such as H, X, and Y shapes, or featuring substantial cavities inside plasmodesmata channels (Faulkner et al. 2008). Unexpectly, complex plasmodesmata did not exhibit an increased size exclusion limit. Primary plasmodesmata, sometimes lacking space between the desmotubule and surrounding plasma membranes, could still facilitate the transport of monomeric GREEN FLUORESCENT PROTEIN (GFP, 27 kDa) (Nicolas et al. 2017; Oparka et al. 1999).

Given the common ancestry of plasmodesmata in land plants, we explored whether the progression of plasmodesmata development in *M. polymorpha* mirrors that of vascular plants. We examined plasmodesmata structures in dormant gemmae, the asexual reproductive units directly obtained from the gemma cup (Fig. 1A, B), under an electron microscope. Hereafter, we will refer to gemmae undergoing development in gemma cups as developing gemmae, well-developed gemmae before germination as dormant gemmae, well-developed gemmae between one to six days after germination as gemmalings, and plants after seven days as thalli (Fig. S1). Our observations revealed plasmodesmata predominantly showing the simple one opening on each side of the cell wall. We noted numerous instances of multiple PDs within a single cell wall throughout the entire section (Fig. 1C). Occasionally, we observed two plasmodesmata aligning nearby (the distance between two plasmodesmata < 200 nm), previously identified as twin plasmodesmata (Fig. 1D, E, white arrowheads) (Faulkner et al. 2008). Additionally, we identified a small number of Y-shaped plasmodesmata in a few cell walls (Fig. 1F). These findings suggest *M. polymorpha* dormant gemmae primarily obtain simple plasmodesmata.

**Fig. 1.**
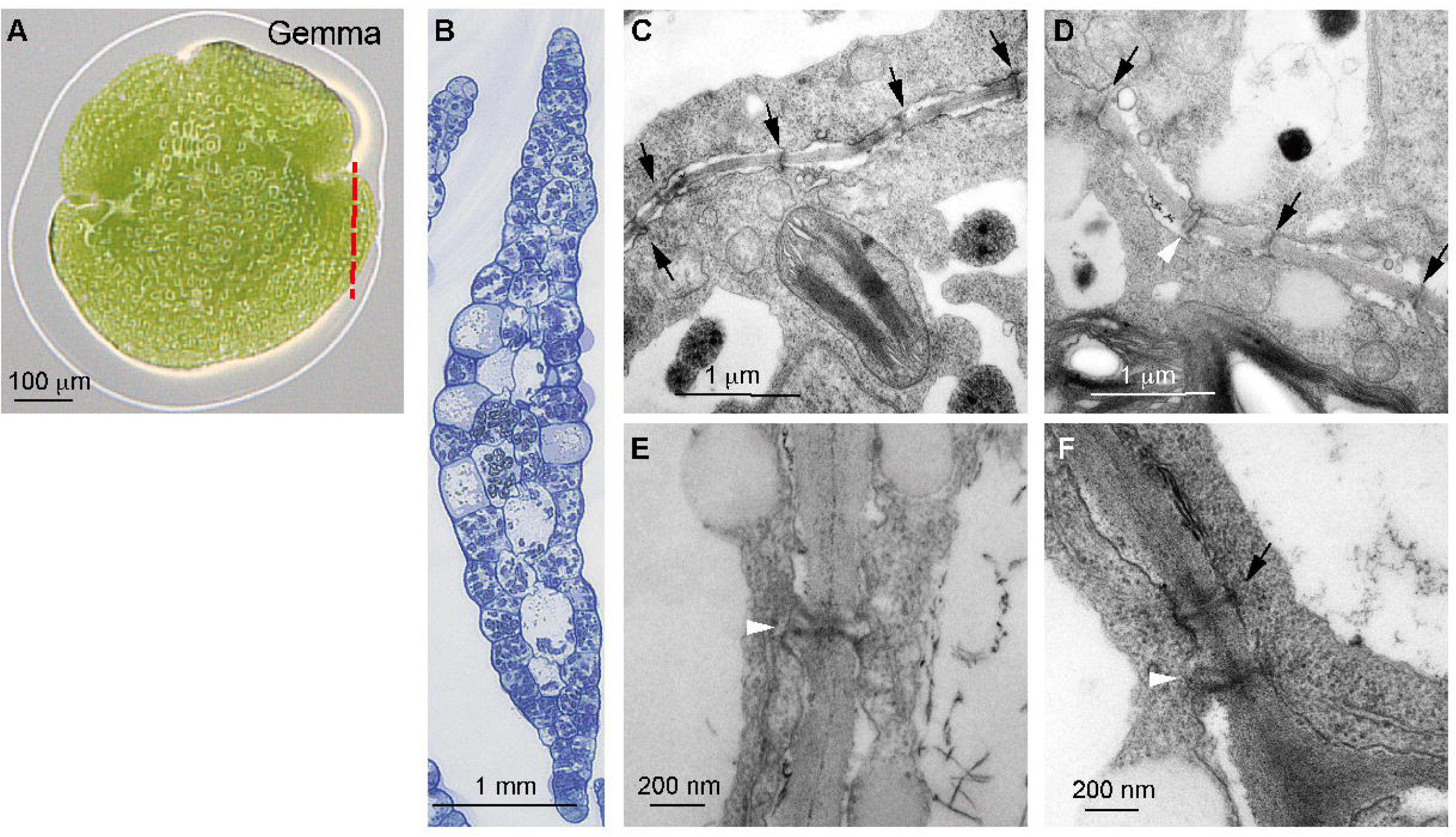
*M. polymorpha* dormant gemmae harbor both simple and complex plasmodesmata. (A) A dormant *M. polymorpha* gemma obtained directly from a gemma cup. The section corresponds to the area indicated by the red dashed line. (B) Overview of the histological section, stained with Toluidine blue. (C) Black arrows indicate plasmodesmata with a single opening on each side of the cell wall. (D) White arrowhead indicates twin plasmodesmata. (E) Twin plasmodesmata at higher magnification. (F) Image of a Y-shaped plasmodesma, indicated by the white arrowhead. Scale bars are provided in each panel.

### Complex plasmodesmata are formed during gemma development and thalli accumulates more complex plasmodesmata

To ascertain whether complex plasmodesmata could be formed during the development of a gemma or generated subsequently after germination, we dissected a gemma cup (Fig. S2 and 2A). We scrutinized the plasmodesmata structure of an early-developing gemma (Fig 2B, the middle black box), a late-developing gemma (Fig. 2B, the top black box), and the mother tissue (the thallus) (Fig. 2B, the bottom black box). Fig. 2S shows the corresponding position of these different gemmae types in the gemma cup. Similar to the dormant gemmae, in the early-developing gemma, most plasmodesmata had one opening on each side of cell walls (Fig. 2C-G, black arrows). However, we also observed twin- (Fig. 2D, E; white arrowheads) and Y-shaped plasmodesmata (Fig. 2F, G; white arrowheads), albeit at a lower frequency. In line with this observation, the late-developing gemma revealed mostly simple and occasional twin plasmodesmata (Fig. 2H, I). Notably, a significant presence of complex plasmodesmata clusters was identified when focusing on the mature thallus (Fig. 2J, K).

**Fig 2.**
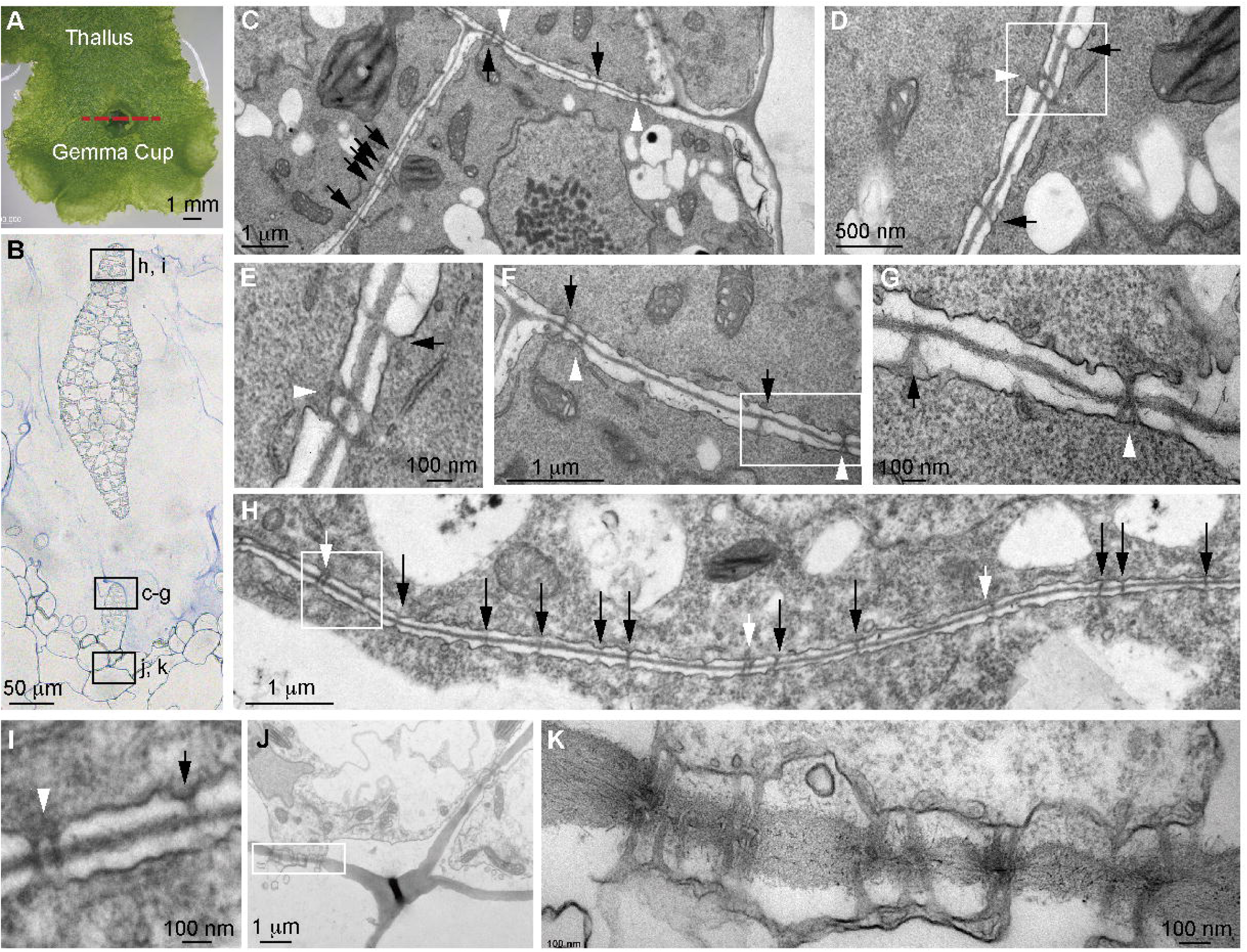
Y-shaped and twin plasmodesmata are present in the developing gemma, while more complex plasmodesmata structures can be observed in the thallus of *M. polymorpha*. (A) A thallus with a gemma cup. The dashed red line marks the dissection site. (B) Overview of the histological section stained with Toluidine blue. Black boxes indicate the developing gemma at different stages (C-G, early; H-I, late) and the thallus tissue (J-K). (C) Plasmodesmata with one opening on each side of the cell wall are indicated by black arrows. White arrowheads mark twin or Y-shaped plasmodesmata. (D) Twin plasmodesmata in the developing gemma. The white box marks the area enlarged in (E). (F) Y-shaped plasmodesmata in the early-developing gemma. The white box marks the area enlarged in (G). (H) Overview of a cross wall in the late-developing gemma. The white box marks the area enlarged in (I). (J) Complex plasmodesmata observed in the mother thallus tissue. The black rod in the center of the image is an artifact from the histological analysis. The white box marks the area enlarged in (K). Scale bars are provided in each panel.

To verify whether the increase in complexity occurs before the maturation of gemmae or after germination, we investigate the plasmodesmata composition of a late-developing gemma and the surrounding thallus cells (Fig. S2F, cyan and purple boxes, respectively). As previously described, we observed majorly simple plasmodesmata with one opening on each side of the cell wall (64%, 402/602 total plasmodesmata in 48 septa of 34 cells). We also calculated the Y-shaped and twin plasmodesmata separately, and we found 28% (178/602) Y-shaped and twin plasmodesmata. We also found 7% (41/602) plasmodesmata are more than three plasmodesmata clustered together in the section (Fig. S3G), indicating that either the insertion of plasmodesmata occurs during the maturation of gemmae, or alternatively, the formation of primary plasmodesmata during the cell wall formation can generate more complex plasmodesmata. In the neighboring thallus cells, we observed 48% (58/120 total plasmodesmata in 22 septa of 14 cells) of simple plasmodesmata, 19% (10/120) of Y-shaped and twin plasmodesmata, and 32% (39/120) clustered plasmodesmata. Among the clustered plasmodesmata, we observed many with enlarged plasmodesmata cavities (Fig. S3H). By using Chi-square analysis, the differences between the two stages have a *p*-value < 0.001, indicating that different stages have significant influences on plasmodesmata complexity. Alongside the altered ratio of intricate structures, we also noted a decrease in plasmodesmata density in the section of mature thallus cell walls (0.24 ± 0.193 PD/μm cell wall, mean ± S.D., N = 11 cells) compared to the late-developing gemma (0.76 ± 0.301 PD/μm cell wall, mean ± S.D., N = 11 cells). The plasmodesmata density differences between the two stages have a *p*-value of 0.0003 (by Student’s *t*-test), indicating significant differences in plasmodesmata density between developing gemmae and thalli (Fig. S3I).

This data suggests that akin to vascular plants, plasmodesmata predominantly adopt a simple structure in early-developing gemmae, while complex plasmodesmata are generated during the maturation of the developing gemma through an unknown secondary plasmodesmata formation mechanism.

### The 26 kDa photoconvertible mEos3.2 fluorescent protein cannot be exported from rhizoid precursor cells

Direct assessment of protein mobility is facilitated by employing photoconvertible or photoinducible fluorescent proteins. In vascular plants and another bryophyte, *P. patens*, monomeric fluorescent proteins have demonstrated intercellular transport through plasmodesmata (Kitagawa and Fujita 2015; Kitagawa et al. 2019). For our study, we opted for the 26 kDa photoconvertible mEos3.2 protein (Mathur et al. 2010), expressed under the 35S promoter known for its ubiquitous and intense expression in *M. polymorpha* (Althoff et al. 2014). We tested the mobility of mEos3.2 in the dormant gemma. We confirmed the ubiquity of p*35S::*mEos3.2 expression, and the green-to-red photoconversion occurred within the first minute of induction (Fig. S4).

To assess the mEos3.2 protein transport, we targeted specific rhizoid precursor cells. Immediately after induction, a sharp increase in red fluorescence was observed in target cells (Fig. 3A, T0). Subsequent tracking of fluorescence changes at ten-minute intervals revealed a lack of red fluorescence increase in neighboring cells within the initial 90 minutes (Fig. 3A). Additionally, both green and red fluorescence exhibited photobleaching with increased scanning repetitions (Fig. 3A and S4). We quantified red and green fluorescence, employing the red-to-green ratio in cells as an indicator of mEos3.2 movement (Fig. 3B, Table S2). A marked increase in the red-to-green ratio in target cells was evident post-induction. In addition, we observed a gradual ratio increase at the induced cell within the 90-minute observation period (Fig. 3B, Table S2). This might be due to the energy of the excitation laser being higher and triggering stronger photobleaching of the green mEos3.2 proteins (Fig. S5). However, none of the surrounding cells exhibited a rise in the red-to-green ratio within the observation period (three representative data points in Fig. 3B; biological repeats are listed in Table S2). To verify whether rhizoid precursor cells are specialized cells so the pleasmodesmata permeability might be differentially regulated, we also performed photo-conversion with other epidermal cells. However, we observed similar patern which no obvious red fluorescence increased during the time observation (Fig S6, Table S3). This outcome implies that the 26 kDa mEos3.2 protein did not undergo outward movement from the induced cells within 90 minutes following photoconversion.

**Fig. 3.**
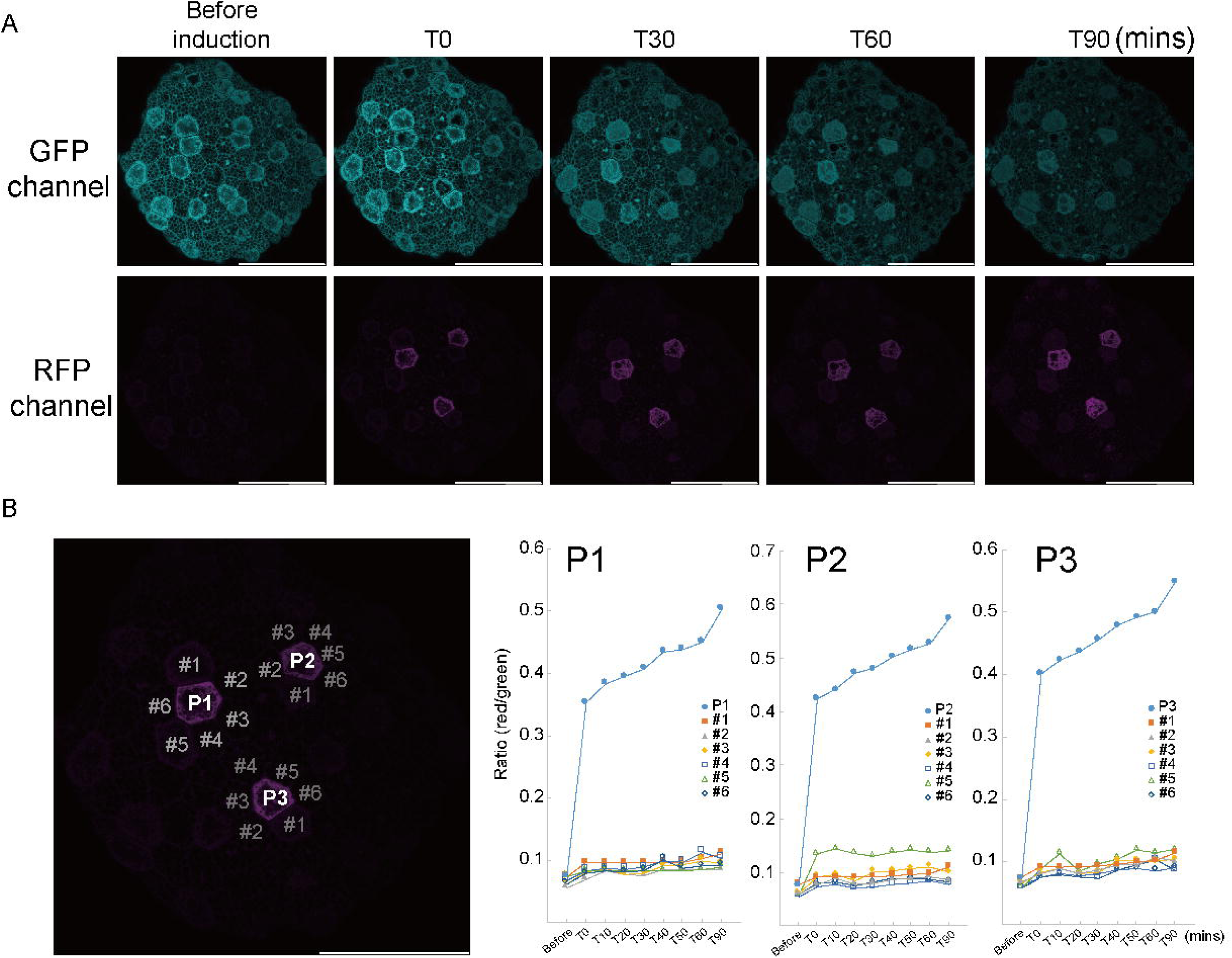
The 26 kDa mEos3.2 protein was not exported to surrounding cells. (A) Three cells were induced for photoconversion, and the fluorescence in both the green and red channels was monitored over 90 minutes. The number following the letter T indicates the time (in minutes) after photoconversion. (B) Charts showing the ratio of GFP to RFP fluorescence in each labeled cell at different time points. The scale bar in each image represents 100 μm.

### Dormant gemmae accumulate callose at plasmodesmata

The immobility of mEos3.2 may indicate a SEL smaller than 26 KDa in *M. polymorpha* dormant gemmae. The regulation of plasmodesmata SEL, often characterized by the accumulation of callose at the neck region of plasmodesmata, is well-documented (De Storme and Geelen 2014; Wu et al. 2018). To investigate callose accumulation in gemmae, we stained with aniline blue, a widely used method for callose visualization in various plant species, including the bryophyte *P. patens* (Huang et al. 2022; Muller et al. 2022). Consistent with the mEos3.2 immobility, bright punctate signals were observed in nearly all cells at the epidermis (Fig. 4A), encompassing cells close to the stem cell niche (Fig. 4B, white arrows). However, weak or no signals were detected in cells next to the apical cells (Fig. 4B, white arrowheads), suggesting potentially a higher PD permeability surrounding the apical cell. At higher magnification, plasmodesmata, as visualized by fluorescent signals, aligned and juxtaposed in adjacent cell walls (Fig. 4C, white arrow). This may indicate that the evolution of callose accumulation at plasmodesmata likely occurred in the common ancestor of land plants. To demonstrate the correlation between callose accumulation and plasmodesmata permeability, a quantification of the staining signals at different domains will provide a good indication. However, due to the hydrophobic surface of *M. polymorpha*, partial or absent staining was frequently observed within the same staining batch (Fig. S7), making the quantification of callose challenging.

**Fig. 4.**
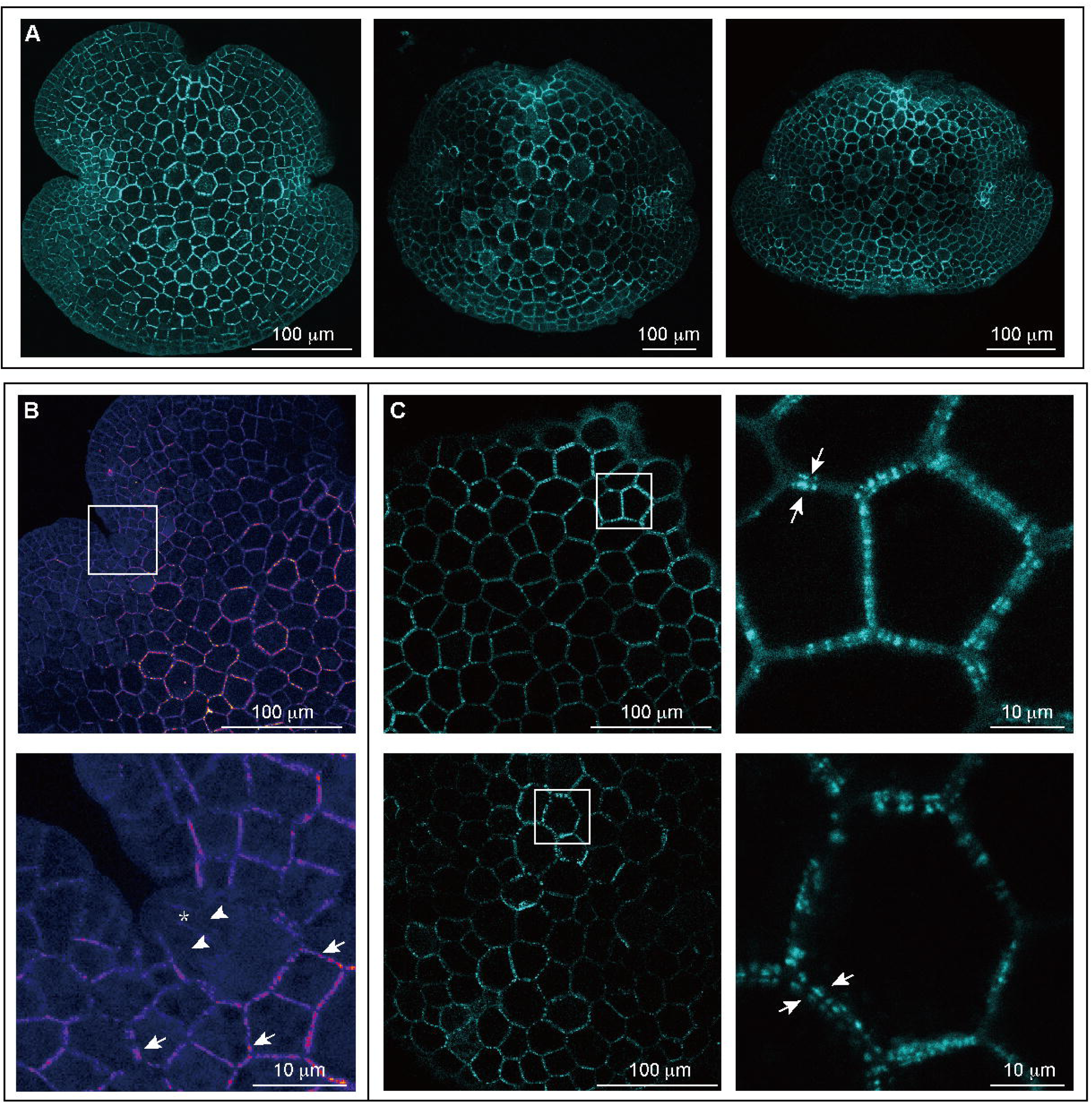
Dormant gemmae accumulate a substantial amount of callose at plasmodesmata. Callose accumulation is visualized using aniline blue staining. (A) Overview of three gemmae. (B) Higher magnification of the apical notch. The white box marks the enlarged area shown below. White arrows indicate the accumulation of callose, while white arrowheads mark cell walls with no apparent callose accumulation. The asterisk indicates the position of the apical cell, located beneath the epidermal cell. (C) Images of epidermal cells in the central region of the gemmae. The figures on the right are enlargements of the areas marked by squares in the left panel. White arrows indicate juxtaposed staining signals across cell walls.

### SYFP2, a 27 kDa fluorescent protein can be transported intercellularly in dormant gemmae and gemmalings

The observed callose accumulation in dormant gemma suggested that the limited permeability of plasmodesmata might hinder mEos3.2 transport. Extending the observation time could counteract this observation. We employed particle bombardment, a method previously demonstrated to transiently express fluorescent proteins in *M. polymorpha* (Westermann et al. 2020). Movement of SYFP2 protein (27 kDa) in *N. benthamiana* leaves was used as a control (Fig. 5A). The spread of the signal shows that SYFP2 moved indeed via plasmodesmata and did not enter symplastic-isolated guard cells (Fig. 5A, asterisk) as their plasmodesmata are degraded after guard cell maturation (Wille and Lucas 1984).

**Fig. 5.**
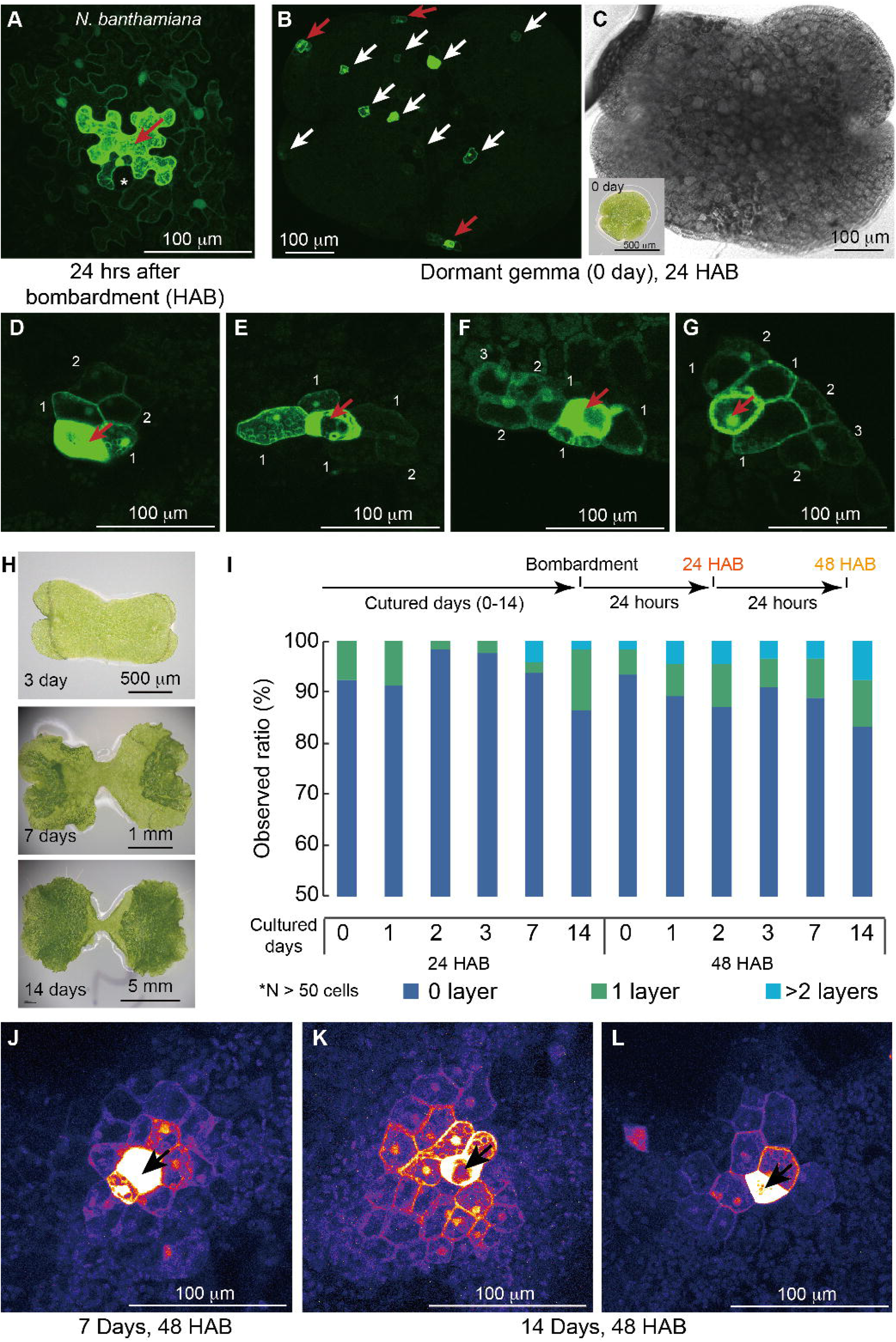
SYFP2 protein can be transported between cells in gemmalings and thalli. (A) Transport of SYFP2 in N. benthamiana leaf epidermal cells. The red arrow marks the bombarded cell. The asterisk indicates the stomata, where guard cells lack plasmodesmata, preventing SYFP2 transport. (B) Representative image of a gemmaling 24 hours after bombardment (HAB). White arrows indicate immobile signals, while red arrows mark mobile events. (C) Bright-field image of (B). The inset shows a dormant gemma (0 days) under a dissecting microscope. (D–G) Representative images of SYFP2 transport in the epidermal cells of a M. polymorpha gemmaling. Numbers in each image correspond to the cell layer receiving SYFP2. (H) Representative images of M. polymorpha at different ages. Dark green tissues in the 7-day and 14-day images are adult tissues, while light green tissues are juvenile tissues. (I) Summary chart of SYFP2 transport in gemmalings and thalli at different ages. (J–L) In adult tissues of the thallus, SYFP2 was transported over a broader area. Black arrows indicate bombarded cells.

Subsequently, *M. polymorpha* dormant gemmae were bombarded with p*35S::SYFP2* plasmids, and expression and movement of SYFP2 proteins were monitored after 24 and 48 hours (HAB) (Fig. 5B, C). SYFP2 expression was evident within 24 HAB, with most signals contained in a single cell (Fig. 5B). However, approximately 10% of bombarded cells exhibited spreading signals outside of the targeted cells, accompanied by a decrease in intensity, resembling observations in *N. benthamiana* leaf epidermis (Fig. 5D-G). Notably, cells receiving mobile SYFP2 were few and appeared to exhibit a random but restricted directionality (Fig. 5D-G), suggesting that SYFP2 can be transported between cells.

While gemmae have been released from dormancy two days after bombardment, it is plausible that the release of plasmodesmata permeability occurs later. Gemmalings cultured on agar medium for 1, 2, and 3 days were bombarded, and SYFP2 transport was monitored (the timeline is illustrated in Fig. 5I). After collecting about 60 bombarded cells from each time point,10-15% of signals spread out within the first 5 days (cultured 3 days plus 48 HAB) (Fig. 5I and Table 1). In general, a higher occurrence of over-two-layer SYFP2 transport was observed after 48 hours than 24 hours (Table 1 and S4).

**Table 1.** The movement of SYFP2 in M. polymorpha epidermis at different ages. The numbers in each cell represent the percentage of events observed at each time point. The bottom row shows the total number of events. The numbers following “D” in the column headers indicate the number of days after incubation on plates, while “24hr” or “48hr” denote the hours after bombardment. In the row headers, “single” refers to fluorescence emission from only the bombarded cell. “1 layer” or “>2 layers” indicate fluorescence emission from one or more than two layers of cells surrounding the bombarded cell, respectively.

|  | D0 |  | D1 |  | D2 |  | D3 |  | D7 |  | D14 |  |
| --- | --- | --- | --- | --- | --- | --- | --- | --- | --- | --- | --- | --- |
| time | 24hr | 48hr | 24hr | 48hr | 24hr | 48hr | 24hr | 48hr | 24hr | 48hr | 24hr | 48hr |
| single | 92.45% | 93.33% | 91.49% | 89.13% | 98.25% | 86.96% | 97.56% | 91.07% | 93.75% | 88.71% | 86.36% | 83.33% |
| 1 layer | 7.55% | 5.00% | 8.51% | 6.52% | 1.75% | 8.70% | 2.44% | 5.36% | 2.08% | 8.06% | 12.12% | 9.09% |
| > 2 layers | 0.00% | 1.67% | 0.00% | 4.35% | 0.00% | 4.35% | 0.00% | 3.57% | 4.17% | 3.23% | 1.52% | 7.58% |
| total | 53 | 60 | 47 | 46 | 57 | 46 | 41 | 56 | 48 | 62 | 66 | 66 |

This result supports the notion that *M. polymorpha* plasmodesmata permeability is consistently stringent at the gemmaling stage, leading to a slow and low transport frequency.

### Extensive movement of the SYFP2 protein can be observed after the transition into thallus

During development, the sink-to-source transition coordinates the plasmodesmata permeability in vascular plants (Oparka et al. 1999). In *M. polymorpha*, a distinct juvenile-to-adult transition is evident between 6 to 7 days after germination (Fig. 5H and S1). The dark green area (adult tissue) forms around the apical notch and gradually becomes the majority of the thallus (Fig. 5H and S1). We were curious whether this stage transition in *M. polymorpha* also regulates plasmodesmata permeability.

Seven- and 14-day-old *M. polymorpha* thalli were subjected to particle bombardment. Even though we observed a slight increase in movement frequency when comparing D14 with D0, we still observed approximately 10-15% of bombarded cells showing spreading fluorescence (Fig. 5I, Table 1, and Table S4), similar to the frequencies observed in other earlier stages. This indicates that plasmodesmata permeability does not change significantly from juvenile to adult stages in *M. polymorpha*. In agreement with the notion, the ANOVA analysis gave a *p*-value of 0.16 for the entire dataset, indicating that the differences in SYFP2 protein mobility across the stages we tested are not statistically significant. We further performed the Tukey HSD analysis to test whether the differences between D0 and D14 are significant. The *p*-value between D0 24hr and D14 24hr is 0.996 while between D0 48hr and D14 48 hr is 0.662, indicating that the differences are not statistically significant. We further calculated the cell numbers that recived the fluorescence from the bombarded cells (Fig. S8 and Table S4), and the statistic analysis remains the same. While there was no significant increase in the movement frequency, we observed a broader transport range in thalli that were either in transition or had reached maturity compared to 0-3 days old gemmalings (Fig 5j-l, comparing to 5d-g). Even though the frequency is low (one event in D7 and two events in D14), this suggests that plasmodesmata permeability in *M. polymorpha* could be regulated when necessary.

### Callose accumulation correlates with the plasmodesmata permeability in *M. polymorpha*

Is plasmodesmata permeability in different areas of a mature thallus correlated with the accumulation of callose? We observed the accumulation of callose in a 14-day-old thallus (Fig. 6a) by aniline blue staining (Fig. 6b-l). The staining is more consistent than what we encountered in gemmalings. We suspect that the formation of air pores largely increased the penetration of aniline blue into the plants (Fig S9). We focused on four different areas: the apical notch, mature tissues, juvenile tissues, and the edge of the thallus. At the apical notch, we rarely detected signals (Fig. 6C, Fig. S9A, B); only when tissues exited the apical notch and started to form air chambers could strong signals be detected (Fig. 6D, Fig. S9A, B). When we focused on mature tissues, we often detected strong punctate signals around cells (Fig. 6E-H, white arrows; Fig. S9D, E). In juvenile tissues and cells along the edge of the thallus, callose accumulation was relatively low compared to cells surrounding air chambers (Fig. 6I-L, Fig. S9E-H). To understand the differences, we quantified the intensity of the signals indifferent in different areas. We measure cells with clear staining surrounding air pores, at the edge, and the juvenile tissue. We obtained an average of 87.4 ± 17.1 a.u./pixel (a.u., arbitrary unit; Mean ± SD, n = 35 cells) at cell walls of the air pore surrounding cells, 75.6 ± 18.9 a.u./pixel (n = 35 cells) at the edge cells, and 58.0 ± 18.0 a.u./pixel (n = 35 cells) of the juvenile cells. The differences between cells surrounding air pores and edge cells are significant, p = 0.004 (Student *t*-test), and the differences between cells surrounding air pores and juvenile cells are also significant, p = 7×10^−10^ (Student *t*-test).

**Fig. 6.**
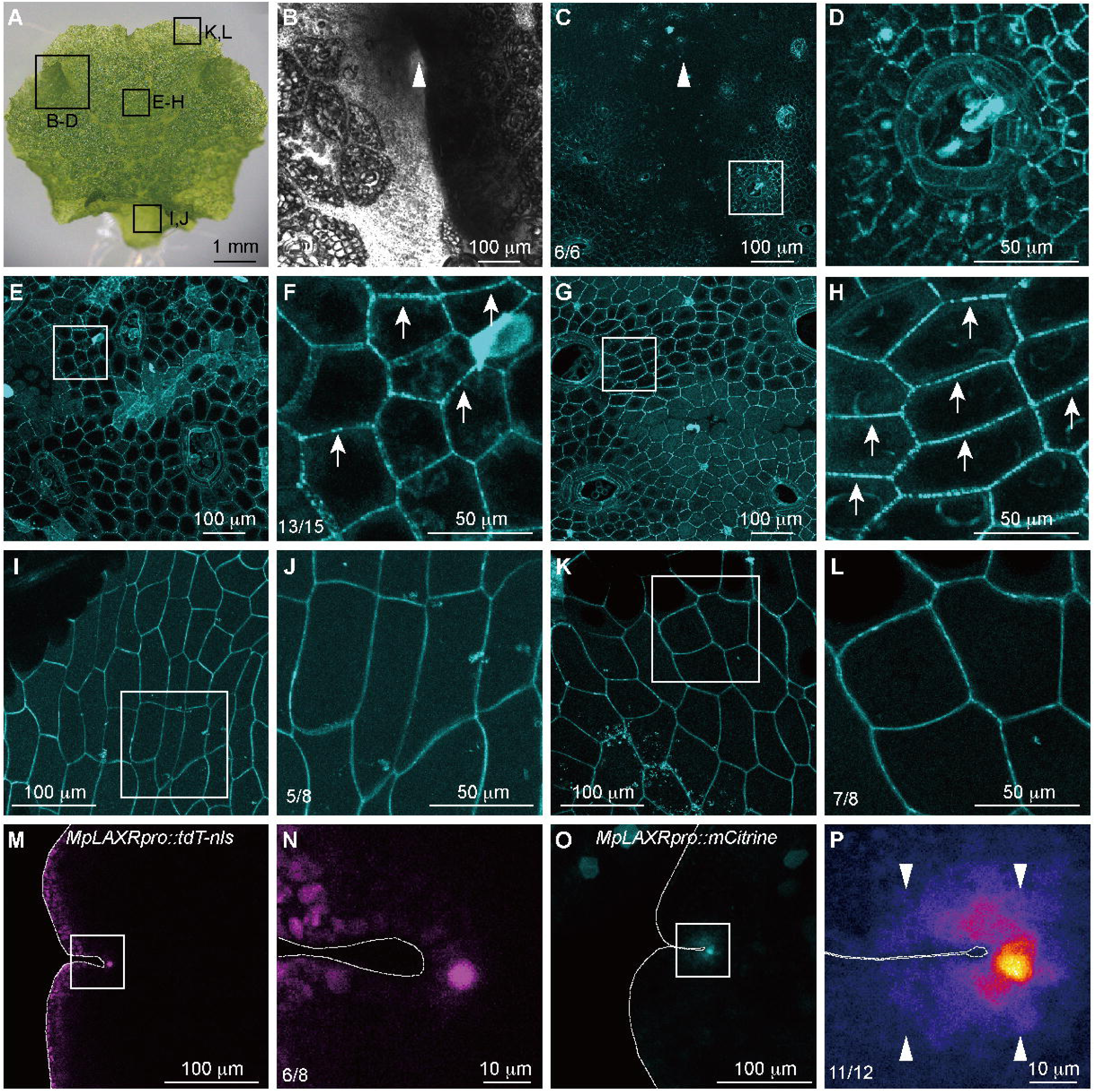
Callose accumulation correlates with the PD permeability in *M. polymorpha*. (A) A 14-day-old thallus. Black boxes mark the areas imaged in the corresponding panels. (B–L) Aniline blue staining images of each selected area. (B) Bright-field image of the apical meristem region. The white arrowhead indicates the apical notch.(C) Aniline blue staining image corresponding to (B). (D) Enlarged area from (C). (E, G) Representative aniline blue staining images of mature epidermal cells. (F, H) Enlarged areas from the white boxes in (E) and (G). White arrows point to punctate signals representing high callose accumulation. (I) Tissues near the edge of the thallus. (J) Little callose accumulation observed in edge tissues. (K) Representative image of juvenile tissues. (L) No clear punctate signals were observed in juvenile epidermal cells. The numbers in the lower-left corners of (C), (F), (J), and (L) indicate the frequencies of similar staining observed across independent repeats. (M, N) Confocal images of MpLAXRpro::tdTOMATO-nls gemmae. (O, P) Confocal images of MpLAXRpro::mCitrine gemmae. The numbers in the lower-left corners of (N) and (P) indicate the frequencies of similar patterns observed across different images. White arrowheads indicate fluorescent signals detected outside the apical cell. Dashed lines outline the shape of the gemma in (M–P).

Finally, we tested if less callose accumulation indicates higher plasmodesmata permeability. To increase the target efficiency, we adopted an apical notch-specific *MpLAXR/MpERF20* promoter that just recently became available, (Ishida et al. 2022; Romani et al. 2023), to express the nucleus-localized tandem TOMATO (tdT-nls, 54 kDa) and the monomeric CITRINE (mCitrine, 27 kDa) fluorescent proteins (Fig. 6M-P). We observed the fluorescence of both transgenic lines in their dormant gemmae. As previously reported, the *MpLAXR::tdT-nls* expressed specifically at the apical cell in two independent lines (Fig. 6M, N). Consistent with the low callose accumulation in the apical notch area (Fig. 4A, B and Fig. 6C, S9A, B), we observed *MpLAXR::*mCitrine had a peak fluorescent intensity at the apical cell and a spray of signals to the surrounding area also in two independent transgenic lines (Fig. 6O, P). Because the apical cell of a dormant gemma will remain inactive, it is less likely the mCitrine protein is distributed to the daughter cells due to cell division. Additionally, under the same observation condition, we did not detect a gradient signal for the tdT-nls protein, suggesting a higher possibility that the fluorescence we detected in *MpLAXR*::mCitrine is from the movement of the mCitrine protein Based on our results, we proposed that plasmodesmata permeability is generally limited, but under different developmental conditions, some cells might escape that limitation and allow higher intercellular transport in both gammae and thalli of *M. polymorph*a.

## Discussion

### The progression of plasmodesmata development is conserved between *M. polymorpha* and vascular plants

Intercellular communication is crucial for multicellular organisms to coordinate their development and growth. Bryophytes and vascular plants diverged approximately 450 million years ago, yet both retain plasmodesmata as the intercellular communication channel, underscoring the indispensability of plasmodesmata for land plants. In vascular plants, primary plasmodesmata, predominantly simple in structure, are known to form during cytokinesis. Subsequently, secondary plasmodesmata, which may exhibit various complex structures, are inserted after cell walls have formed (Faulkner et al. 2008). Previous research on the liverwort *Monoclea gottschei* demonstrated the formation of both simple and complex (branched) plasmodesmata (Cook et al. 1997). Similarly, in *M. polymorpha*, our observations revealed predominantly simple PD with few twin- or Y-shaped structures and occasionally more than three plasmodesmata clustered in developing gemmae, a characteristic retained in dormant gemmae. Additionally, in thalli, the number of plasmodesmata in a cell wall section decreased (Fig. S1). However, more complex plasmodesmata structures became evident (Fig. 2J, S1), suggesting the presence of a secondary mechanism in *M. polymorpha*, possibly involved in either modifying pre-existing plasmodesmata or forming new ones. It is also worth noting that in our research, we did not separate the effect between different tissues in the thalli and we compared the overall observation between the developing gemmae and the cells surrounding the gemmae cup (Fig 2 and S1). Therefore, we propose that the formation of more than three plasmodesmata occurs during the development of gemmae; in thalli, this process continues, resulting in increasing the percentage and enlarging cavities of complex plasmodesmata (Fig S1). Similar to our results, Wegner and Ehlers also observed a decrease in plasmodesmata density and an increase in plasmodesmata complexity along the development. The increase in complexity could be attributed to the incomplete plasmodesmata-twinning process (Wegner and Ehlers 2024). Together, our results indicate that the progression of plasmodesmata development and the mechanisms of plasmodesmata formation may have evolved in the most common ancestor of land plants.

### The accumulation of callose around the plasmodesmata region is evolutionarily conserved

The presence of callose is a general plasmodesmata regulatory mechanism in vascular plants. Many developmental processes have been indicated to regulate callose accumulation to perform symplasmic isolation, such as the establishment of dormancy, lateral root initiation, and stress responses (De Storme and Geelen 2014; Wu et al. 2018). In addition, one of the charophytic algae, *Chara coralline*, which obtains structures analogous to PD with an enclosed membrane structure instead of the internal ER at the center of the structure, also accumulates callose at the plasmodesmata analogous areas, suggesting a potential evolutionary conserved mechanism to deposit callose around plasmodesmata (Faulkner et al. 2009). It is not difficult to picture that the most common ancestor of land plants has already obtained this mechanism, and both bryophytes and vascular plants inherited it. Indeed, we observed strong punctate signals located at the periphery of cells in dormant gemmae (Fig. 4). We also observed that there is low callose accumulation on cell walls surrounding the apical notch region (Fig. 4B, 6C, and S9A, B), suggesting that a high symplastic connection might play a role for the development/maintenance of the apical notch in *M. polymorpha*. Consistent with the callose accumulation, we also observed the transport of mCitrine into cells surrounding the apical cell (Fig. 6O, P). In vascular plants, it has been shown that there is a domain at the shoot apical meristem where fluorescent-labeled Dextrans can freely diffuse (Rinne and van der Schoot 1998). In the root apical meristem area, monomeric GFP can also diffuse through plasmodesmata, even though there seems to be no space between the plasma membrane and the desmotubule (Nicolas et al. 2017). Combining our data with the known evidence from vascular plants, we propose that the high permeability of the apical meristem might be a common characteristic of the meristematic domain that evolved before the divergence of bryophytes and vascular plants.

### The plasmodesmata permeability is generally stringent in *M. polymorpha* but might be regulated in specific tissues

In vascular plants, the activity of plasmodesmata is tightly regulated and is crucial for many developmental events, such as entering dormancy or forming lateral roots (Benitez-Alfonso et al. 2013; De Storme and Geelen 2014). *M. polymorpha* gemmae are formed in gemma cups and remain dormant until the imbibition of moisture (Shimamura 2015). The highly accumulated callose around plasmodesmata in dormant gemmae suggests that the regulation of dormancy in *M. polymorpha* may correlate with the permeability of plasmodesmata as well (Fig. 4 and Fig. 5). However, the frequency of SYFP2 movement in the first 5 days after imbibition shows no significant increase (Fig. 3, Fig. 5, and Table 1). This data suggests that the permeability of plasmodesmata and gemmae dormancy regulation might be independent mechanisms in *M. polymorpha*.

Sink (young) tissues generally have more simple plasmodesmata with larger SEL to support the effective exchange of nutrients and molecules in vascular plants (Nicolas et al. 2017; Oparka et al. 1999). In *M. polymorpha*, there is no obvious sink-to-source transition but a clear juvenile-to-adult transition (Shimamura 2015). We expected that similar to vascular plants, with the increase of plasmodesmata complexity, the permeability would be higher in gemmalings and would decrease after transitioning to thalli. Instead, we found a comparable movement frequency in both gemmalings and thalli, indicating a generally stringent regulation of plasmodesmata permeability in *M. polymorpha*. Consistent with this notion, a high accumulation of callose was found in most of the investigated tissues, except for the apical notches, edge cells, and juvenile tissues at thalli (Fig. 4, 6C-L, and S9). Together with the increase in transport range in the adult thallus (Fig. 5J-L, S8 and Table S4), our results indicate that PD permeability in *M. polymorpha* might be regulated by developmental cues.

In summary, our research provides fundamental insights into the properties of plasmodesmata in *M. polymorpha*. With its simple genome, *M. polymorpha* has the potential to serve as a model system for studying the conserved composition and function of plasmodesmata. Several aspects warrant further investigation, such as whether callose synthases are involved in the developmental regulation of *M. polymorpha* or whether *M. polymorpha* and Arabidopsis share a conserved plasmodesmata composition. Exploring these questions will enhance our understanding of the critical roles plasmodesmata play in plant development and their evolutionary significance in land plants.

## Acknowledgments

This work was financially supported by the Columbus program of National Science and Technology Council (NSTC) (NSTC 112-1636-B-005-001-), the Advanced Plant and Food Crop Biotechnology Center from The Featured Areas Research Center Program within the framework of the Higher Education Sprout Project, and Yushan Young Fellow program by the Ministry of Education (MOE) (MOE-109-YSFAG-0006-001-P1) in Taiwan. We gratefully thank Prof. Takayuki Kohchi, Prof. Ryuichi Nishihama, and Prof. Dolf Weijers for the support of plasmids and materials. We thank the live-cell-imaging core lab and the electron microscope division, the cell biology core lab in the Institute of Plant and Microbial Biology, Academia Sinica, for all the microscopy support. We also thank Dr. Barbara Kloeckener Gruissem for the critical review and editing of the manuscript.

## Data availability

The data underlying this article are available in the article and its online supplementary material.

## Author contributions

KJL planned and designed the research. JYH, JHH, HYC, and KJL performed experiments and analyzed data. KJL wrote the manuscript. JYH, JHH, and HYC contributed equally

