## Supplementary material for "Exploring The Formation And Permeability Of Plasmodesmata In The Liverwort, *Marchantia polymorpha*": Fig. S

Supplemntal Figure S1, Hsu et al.

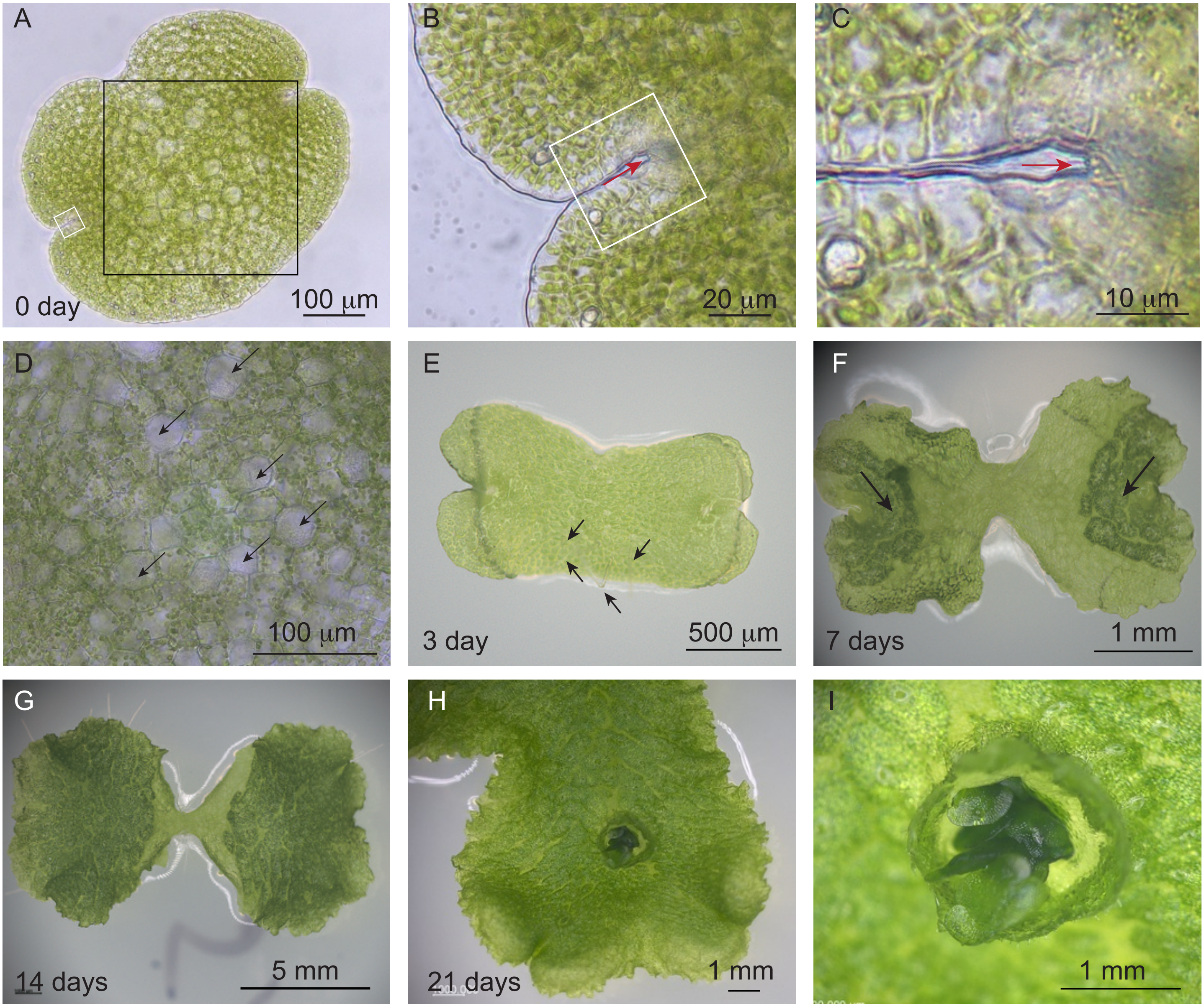


**Fig. S1. Illustrations of *M. polymorpha* at different developmental stages.**
(A) A dormant gemma. The white box indicates the apical notch, which is enlarged in (B) and (C). The black square is enlarged in (D). (B) The apical notch. The white box is further enlarged in (C). The red arrow indicates the location of the apical cell. (C) Enlarged view of the apical notch. The red arrow points to the apical cell located beneath the epidermal cells. (D) The apical cells at the center of the gemma. Black arrows indicate the rhizoid precursor cells, which contain no chloroplasts. (E) A three-day-old gemmaling. Black arrows indicate the protruding rhizoids. (F) A seven-day-old thallus. Black arrows indicate the dark green, mature tissues. (G) A fourteen-day-old thallus. Most of the tissues are mature. (H) A mature thallus with one gemma cup. (I) Enlarged view of the gemma cup in (H), showing many gemmae being generated and pushed out from the gemma cup. Scale bars are as indicated in each figure.

Supplemntal Figure S2, Hsu et al.

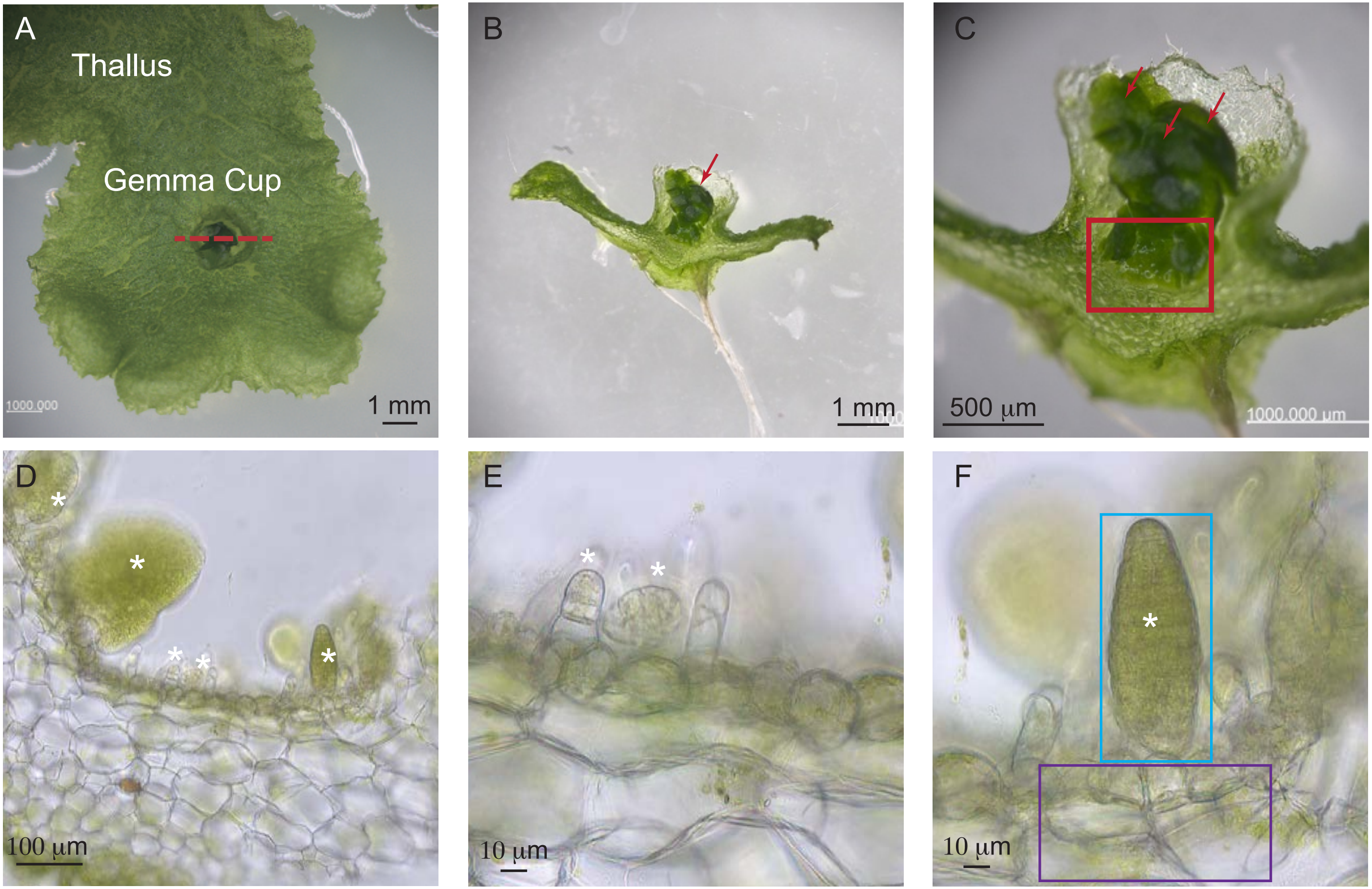


**Fig. S2. Illustration of tissue sections for electron microscopy.**
(A) Image of a mature thallus with one gemma cup. The red dashed line indicates the position of the final section. (B, C) Side views of the vertically sectioned tissue. Red arrows indicate dormant gemmae. (D–F) Following the removal of most mature gemmae, many developing gemmae are observed at the bottom of the gemma cup. (E, F) Magnified views showing a few early-developing gemmae (E) and a late-developing gemma (F). The cyan and purple boxes highlight the late-developing gemma and the mother tissue analyzed for plasmodesmata density, respectively. White asterisks indicate developing gemmae. Scale bars are as indicated in each figure.

### Supplemntal Figure S3, Hsu et al.

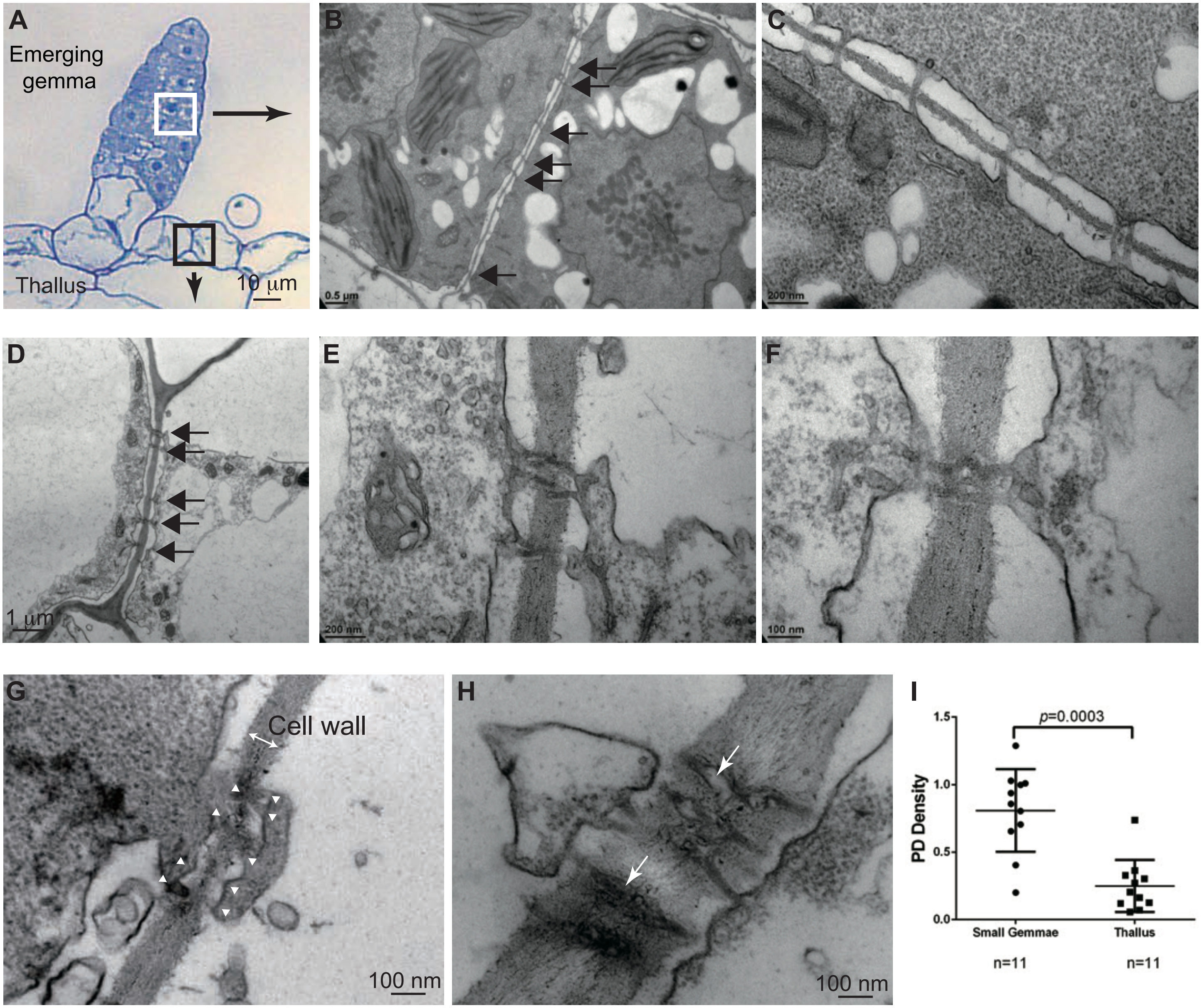


**Figure S3. The strcuture and density of plasmodesmata differ in developing gemma and mature thallus.**

(A) A developing gemma. (B, C) In early-developing gemmae, simple plasmodesmata were predominantly observed, with twin plasmodesmata occurring at low frequency. (D–F) In thalli, more complex plasmodesmata were frequently observed. Plasmodesmata are indicated by black arrows. (G) In late-developing gemmae, complex plasmodesmata were observed at low frequency. White arrow heads indicate the apertures of the plasmodesmata. (H) In thalli, complex plasmodesmata often exhibit enlarged cavities, as indicated by white arrows. (I) The average plasmodesmata density in developing gemmae and mature thalli. Each dot and square represent data from a single cell. The upper and lower horizontal lines correspond to the third and first quartiles, respectively, with the middle line representing the median. The *p*-value was calculated using Student's *t*-test.

### Supplemental Fig. S4, Hsu et al.

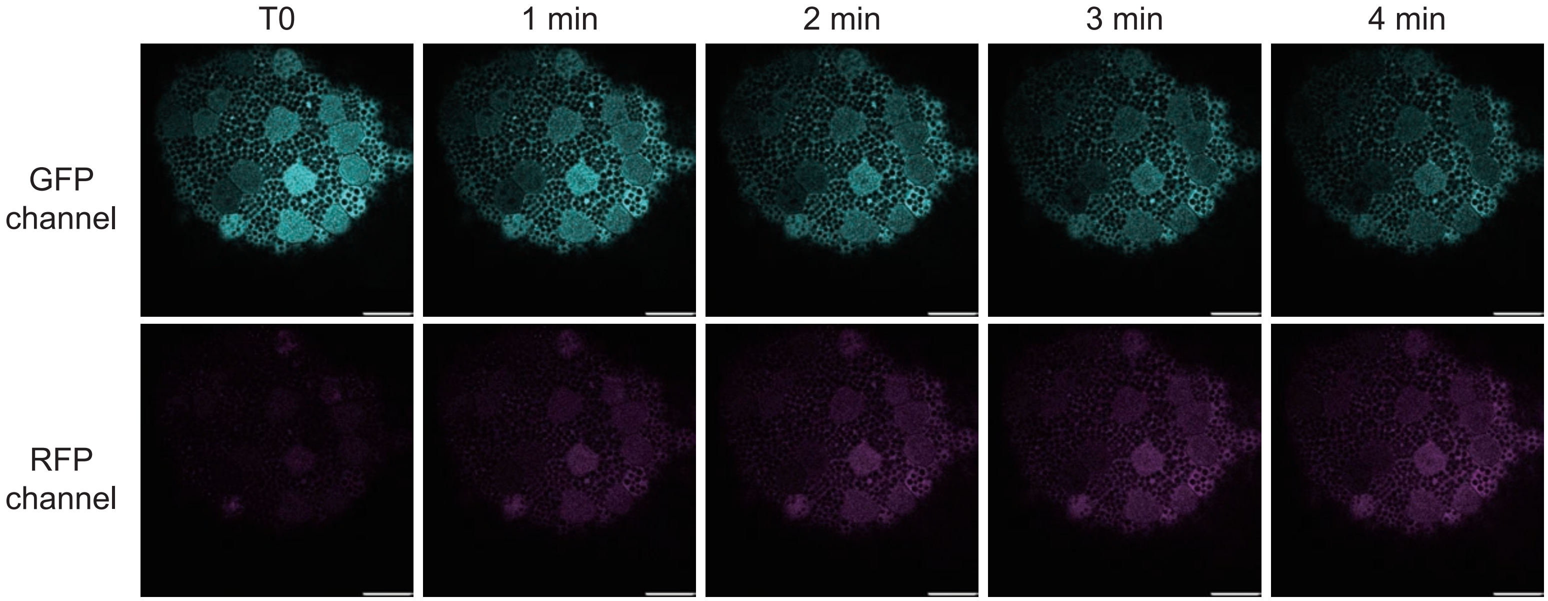


**Fig. S4 The expression and photoconversion of p*35S::mEos3.2* transgenic line.**
The gemma of p*35S::mEos3.2* transgenic *M. polymorpha* expressed the mEos3.2 ubiquitously. The photoconversion was induced by continuous scanning with 405 nm UV and imaging at the same time. Weak to no RFP signals was detected at T0 and the conversion from green to red were clearly observed as the scanning time increased. The scale bar in each figure indicates 50 μm.

### Supplemental Fig. S5, Hsu et al.

A

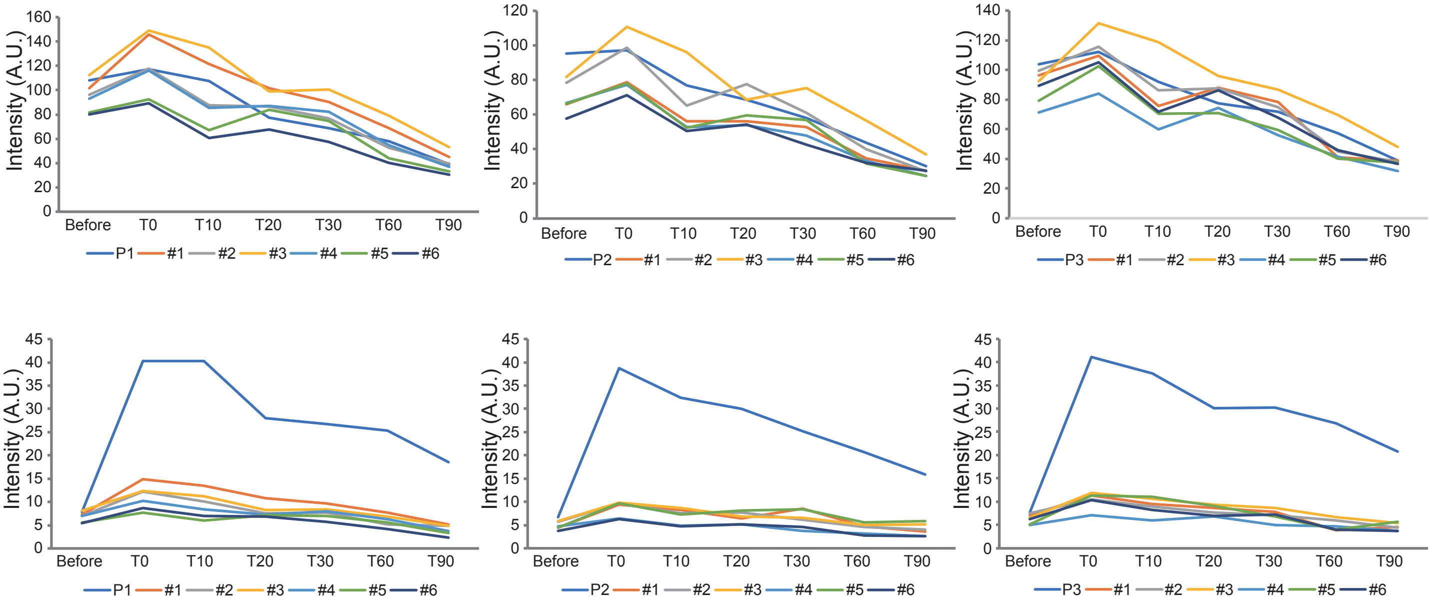


B

**The green channel fluorescence intensity (A.U.)**

| | P1 | #1 | #2 | #3 | #4 | #5 | #6 |
|---|---|---|---|---|---|---|---|
| B4 | 107.9709499 | 101.5806265 | 96.0741263 | 112.4621133 | 92.68724778 | 81.76570145 | 80.10214542 |
| T0 | 117.0835587 | 145.8161202 | 117.596316 | 148.9258078 | 115.9105736 | 92.29209763 | 89.41790351 |
| T10 | 107.5315148 | 121.6599352 | 87.34417434 | 135.1999521 | 85.31385039 | 67.26893799 | 60.48111255 |
| T20 | 77.43125353 | 101.4251734 | 86.61377658 | 99.05542968 | 86.9196668 | 83.80319415 | 67.79604258 |
| T30 | 68.74310913 | 90.1158581 | 77.02762151 | 100.4305688 | 82.10019315 | 74.46564682 | 57.27499258 |
| T60 | 57.686 | 68.854 | 52.637 | 78.935 | 54.429 | 44.078 | 40.250 |
| T90 | 39.024 | 45.026 | 39.528 | 52.817 | 36.662 | 32.908 | 30.457 |
| Intensity % compare to B4 | 36.143 | 44.325 | 41.144 | 46.964 | 39.555 | 40.247 | 38.022 |

| | P2 | #1 | #2 | #3 | #4 | #5 | #6 |
|---|---|---|---|---|---|---|---|
| B4 | 95.05733232 | 65.97766732 | 78.27152927 | 81.71444394 | 66.56501199 | 66.10272921 | 57.66419201 |
| T0 | 97.08458527 | 78.67481459 | 98.63782636 | 110.5855688 | 77.19205439 | 78.0092282 | 70.9268305 |
| T10 | 76.66474892 | 55.99869295 | 65.01526032 | 95.99784842 | 52.72032402 | 52.42105987 | 50.30798646 |
| T20 | 68.50539127 | 56.17193762 | 77.52606932 | 68.55307708 | 53.91947306 | 59.59981478 | 54.29285543 |
| T30 | 58.04871862 | 52.81543525 | 61.06496353 | 75.27345674 | 47.68216688 | 56.91727323 | 42.69882776 |
| T60 | 43.536 | 34.470 | 39.816 | 56.346 | 32.933 | 31.481 | 31.964 |
| T90 | 30.270 | 27.535 | 27.225 | 36.722 | 24.475 | 24.562 | 27.312 |
| Intensity % compare to B4 | 31.84342387 | 41.73383949 | 34.78267762 | 44.9397756 | 36.76817489 | 37.15700737 | 47.36319411 |

| | P3 | #1 | #2 | #3 | #4 | #5 | #6 |
|---|---|---|---|---|---|---|---|
| B4 | 103.6321186 | 96.57359738 | 99.32315585 | 92.30484346 | 71.34169724 | 79.09224241 | 89.1331136 |
| T0 | 112.1292373 | 109.4191727 | 115.4500211 | 131.5590752 | 84.10147955 | 102.4932345 | 105.0536363 |
| T10 | 92.00016869 | 75.92607828 | 86.2654938 | 118.9430283 | 59.92836937 | 70.26537074 | 71.88700638 |
| T20 | 77.29094971 | 87.96803657 | 87.51697064 | 95.86163431 | 74.41221547 | 71.06626402 | 86.30523083 |
| T30 | 71.64310293 | 78.19940933 | 74.64480271 | 86.78553197 | 55.87723605 | 59.43897031 | 67.78896959 |
| T60 | 57.094 | 41.127 | 45.205 | 69.527 | 41.466 | 40.320 | 46.075 |
| T90 | 38.943 | 38.752 | 38.169 | 48.056 | 31.663 | 37.649 | 36.536 |
| Intensity % compare to B4 | 37.57857821 | 40.1270479 | 38.4294296 | 52.06242137 | 44.38282525 | 47.60190329 | 40.99009297 |

**The red channel fluorescence intensity (A.U.)**

| | P1 | #1 | #2 | #3 | #4 | #5 | 6% |
|---|---|---|---|---|---|---|---|
| B4 | 7.847974478 | 7.284146971 | 6.926043383 | 8.177799701 | 6.954319263 | 5.631871437 | 5.410793732 |
| T0 | 40.26147797 | 14.95610787 | 12.21647191 | 12.38725103 | 10.21832222 | 7.680980197 | 8.625248923 |
| T10 | 40.31917847 | 13.49534253 | 10.07007037 | 11.25506411 | 8.447981092 | 6.014826339 | 7.025246707 |
| T20 | 28.06553607 | 10.77042557 | 7.566656566 | 8.266123261 | 7.253565496 | 7.09858863 | 6.803941018 |
| T30 | 26.68714828 | 9.662717875 | 7.712241956 | 8.461663394 | 7.966911274 | 6.982757805 | 5.66570659 |
| T60 | 25.303 | 7.658 | 5.135 | 6.830 | 6.313 | 5.522 | 4.179 |
| T90 | 18.579 | 5.209 | 4.819 | 4.904 | 3.698 | 3.393 | 2.395 |
| Intensity % compare to T0 | 46.147 | 34.830 | 39.445 | 39.587 | 36.190 | 44.174 | 27.771 |

| | P2 | #1 | #2 | #3 | #4 | #5 | #6 |
|---|---|---|---|---|---|---|---|
| B4 | 6.736142704 | 4.415240861 | 5.708131125 | 5.844247501 | 4.705276774 | 4.424102423 | 3.781736668 |
| T0 | 38.71269795 | 9.362220928 | 9.73398722 | 9.820847246 | 6.381531487 | 9.690987737 | 6.340952631 |
| T10 | 32.34999204 | 8.312474233 | 7.565103386 | 8.733319105 | 4.946985699 | 7.312203727 | 4.749630581 |
| T20 | 30.01747906 | 6.423718108 | 7.775932962 | 6.841284055 | 5.215073913 | 8.088744796 | 5.106197164 |
| T30 | 25.24800838 | 8.603020996 | 6.180871013 | 6.519926995 | 3.783061814 | 8.47810184 | 4.599046021 |
| T60 | 20.671 | 4.741 | 4.620 | 5.078 | 3.256 | 5.554 | 2.764 |
| T90 | 15.950 | 3.601 | 4.083 | 5.188 | 2.641 | 5.856 | 2.596 |
| Intensity % compare to T0 | 41.201 | 38.461 | 41.943 | 52.823 | 41.390 | 60.430 | 40.941 |

| | P3 | #1 | #2 | #3 | #4 | #5 | #6 |
|---|---|---|---|---|---|---|---|
| B4 | 7.594884627 | 7.049793156 | 7.444548028 | 6.495207867 | 4.930883826 | 5.086794509 | 6.207822779 |
| T0 | 41.07303547 | 11.33370173 | 10.4447607 | 11.84431206 | 7.08096767 | 11.36871515 | 10.28318703 |
| T10 | 37.6059434 | 9.48732502 | 8.935764108 | 10.53783903 | 5.915976476 | 11.04669321 | 8.173260787 |
| T20 | 30.01697247 | 8.664578748 | 7.670194133 | 9.320982506 | 6.838672379 | 9.096312536 | 6.985350682 |
| T30 | 30.17571025 | 7.732010323 | 7.095906311 | 8.616324992 | 4.956840994 | 6.763074095 | 7.163155336 |
| T60 | 26.869 | 3.856 | 5.953 | 6.694 | 4.671 | 4.031 | 4.000 |
| T90 | 20.780 | 4.521 | 4.356 | 5.397 | 3.738 | 5.621 | 3.654 |
| Intensity % compare to T0 | 50.59264804 | 39.89257379 | 41.70795539 | 45.56234735 | 52.78663066 | 49.43964752 | 35.53324211 |

**Fig. S5. Quantification of mEos3.2 photo-conversion in the green and red channels.**
(A) Intensity line charts for each position: the upper chart represents data from the green channel, while the lower chart shows data from the red channel. (B) The raw data corresponding to each chart. A.U., arbitrary units.

### Supplemental Fig. S6, Hsu et al.

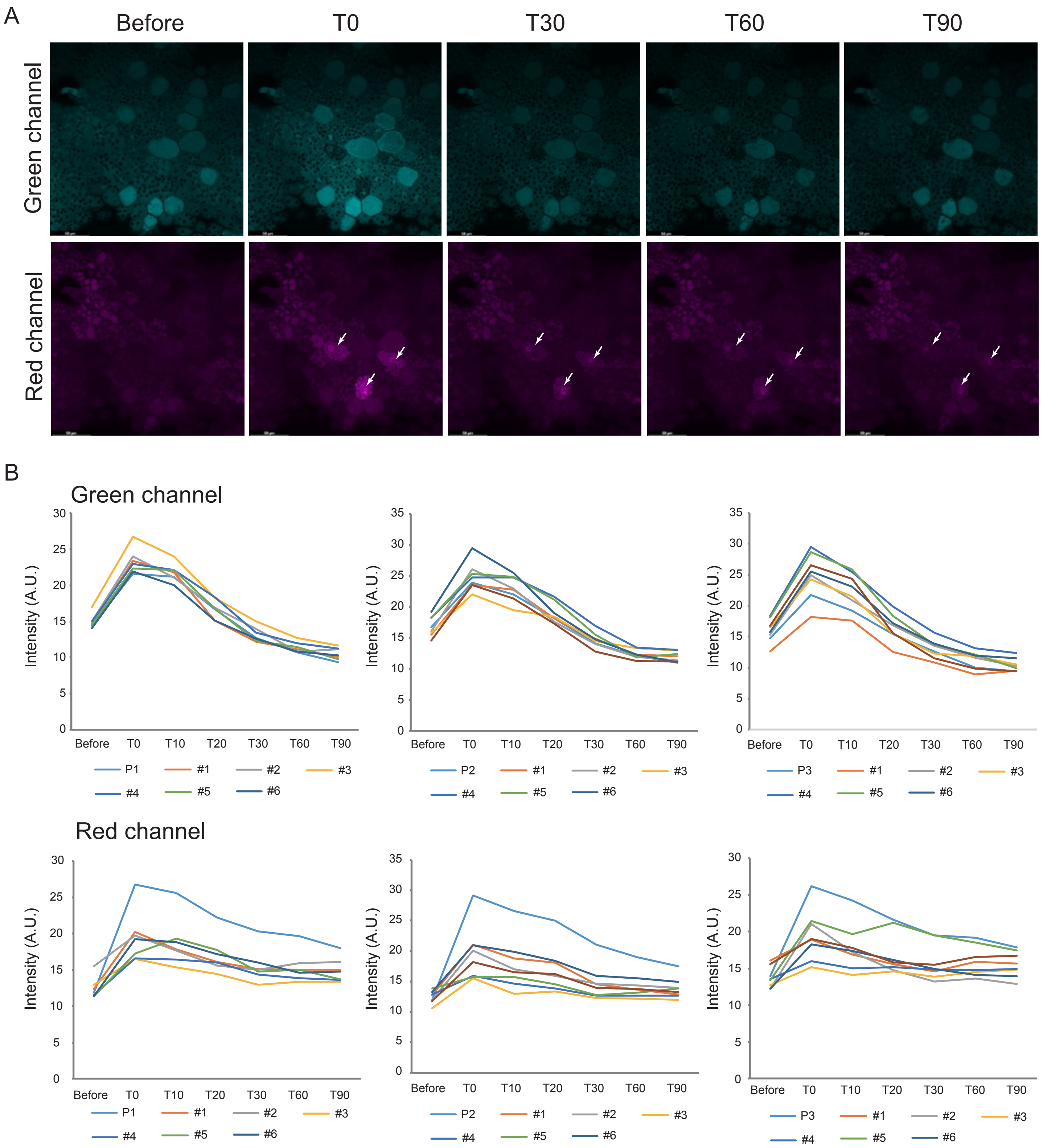


**Fig. S6. mEos3.2 photo-conversion in epidermal cells.**
(A) Representative images of photo-conversion in three epidermal cells. White arrows indicate the cells targeted for conversion. Scale bars are as indicated in each figure. (B) Intensity line charts for each position: the upper chart represents data from the green channel, while the lower chart represents data from the red channel. A.U., arbitrary units.

### Supplemental Fig. S7, Hsu et al.

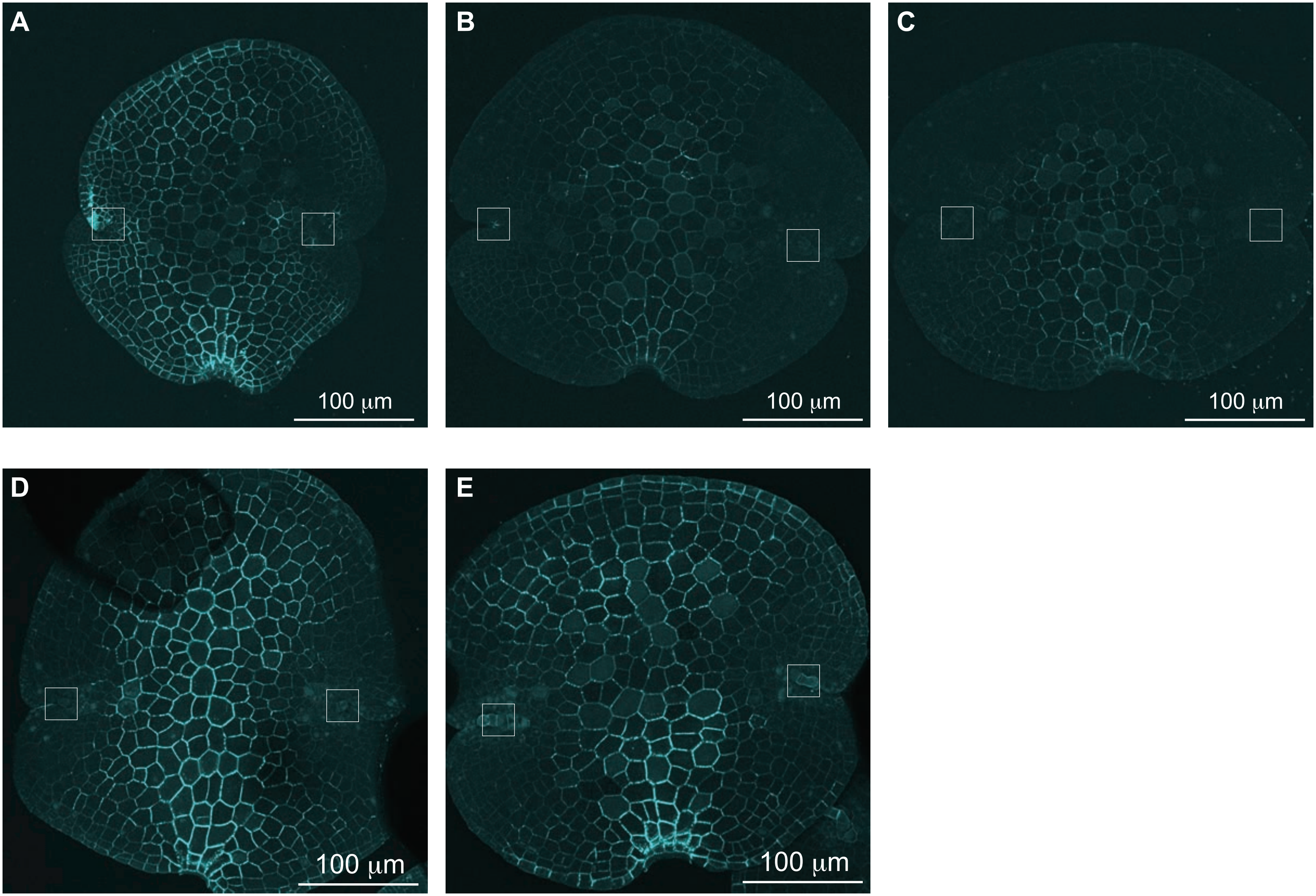


**Fig S7. The limitations of aniline blue staining with *M. polymorpha* dormant gemmae.**
The surface of *M. polymorpha* is highly hydrophobic, leading to uneven staining. In the same batch of samples, uneven staining was frequently observed across different gemmae. (A) The center, upper right, and right apical notch areas show dimmer signals. (B, C) Only the center and lower areas are stained. (D, E) The most common staining pattern is well-stained centers with no signal near the apical notches. The white boxes mark the apical notch area in each figure. Note that in (D) and (E), the apical notches are covered by mucilage cells.

### Supplemental Fig. S8,Hsu et al

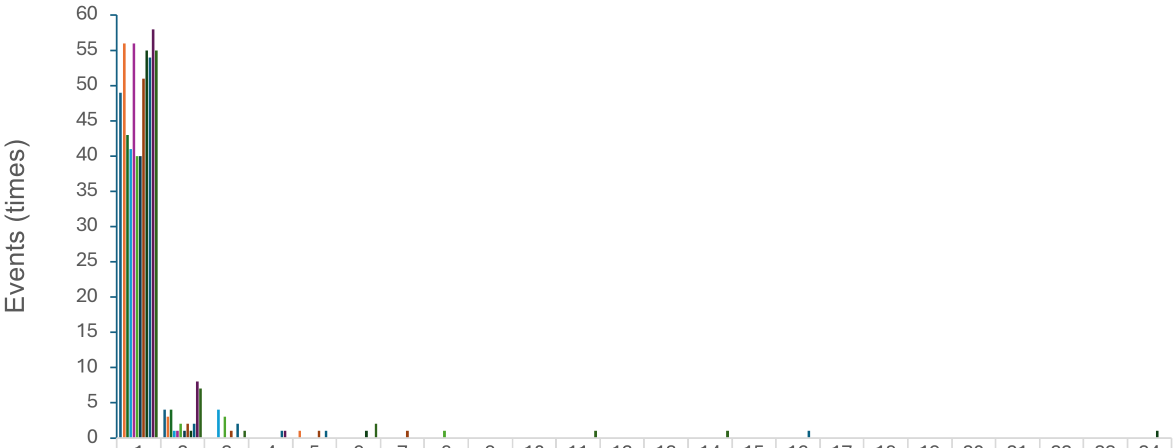


| | 1 | 2 | 3 | 4 | 5 | 6 | 7 | 8 | 9 | 10 | 11 | 12 | 13 | 14 | 15 | 16 | 17 | 18 | 19 | 20 | 21 | 22 | 23 | 24 |
|---|---|---|---|---|---|---|---|---|---|---|---|---|---|---|---|---|---|---|---|---|---|---|---|---|
| D0 24hr | 49 | 4 | | | | | | | | | | | | | | | | | | | | | | |
| D0 48hr | 56 | 3 | | | 1 | | | | | | | | | | | | | | | | | | | |
| D1 24hr | 43 | 4 | | | | | | | | | | | | | | | | | | | | | | |
| D1 48hr | 41 | 1 | 4 | | | | | | | | | | | | | | | | | | | | | |
| D2 24hr | 56 | 1 | | | | | | | | | | | | | | | | | | | | | | |
| D2 48hr | 40 | 2 | 3 | | | | | 1 | | | | | | | | | | | | | | | | |
| D3 24hr | 40 | 1 | | | | | | | | | | | | | | | | | | | | | | |
| D3 48hr | 51 | 2 | 1 | | 1 | | 1 | | | | | | | | | | | | | | | | | |
| D7 24hr | 55 | 1 | | | | 1 | | | | | | | | | | | | | | | | | | 1 |
| D7 48hr | 54 | 2 | 2 | 1 | 1 | | | | | | | | | | | 1 | | | | | | | | |
| D14 24hr | 58 | 8 | | 1 | | | | | | | | | | | | | | | | | | | | |
| D14 48hr | 55 | 7 | 1 | | | 2 | | | | | 1 | | | 1 | | | | | | | | | | |

Cells with fluorescence

**Fig S8. Cell numbers receiving SYFP2 fluorescence from bombarded cells.**
The X-axis indicates the numbers of cells have fluorescence and the Y-axis indicates the event numbers in each experiment setup. The data is listed in the spreadsheet below the bar chart.

### Supplemental Fig. S9, Hsu et al.

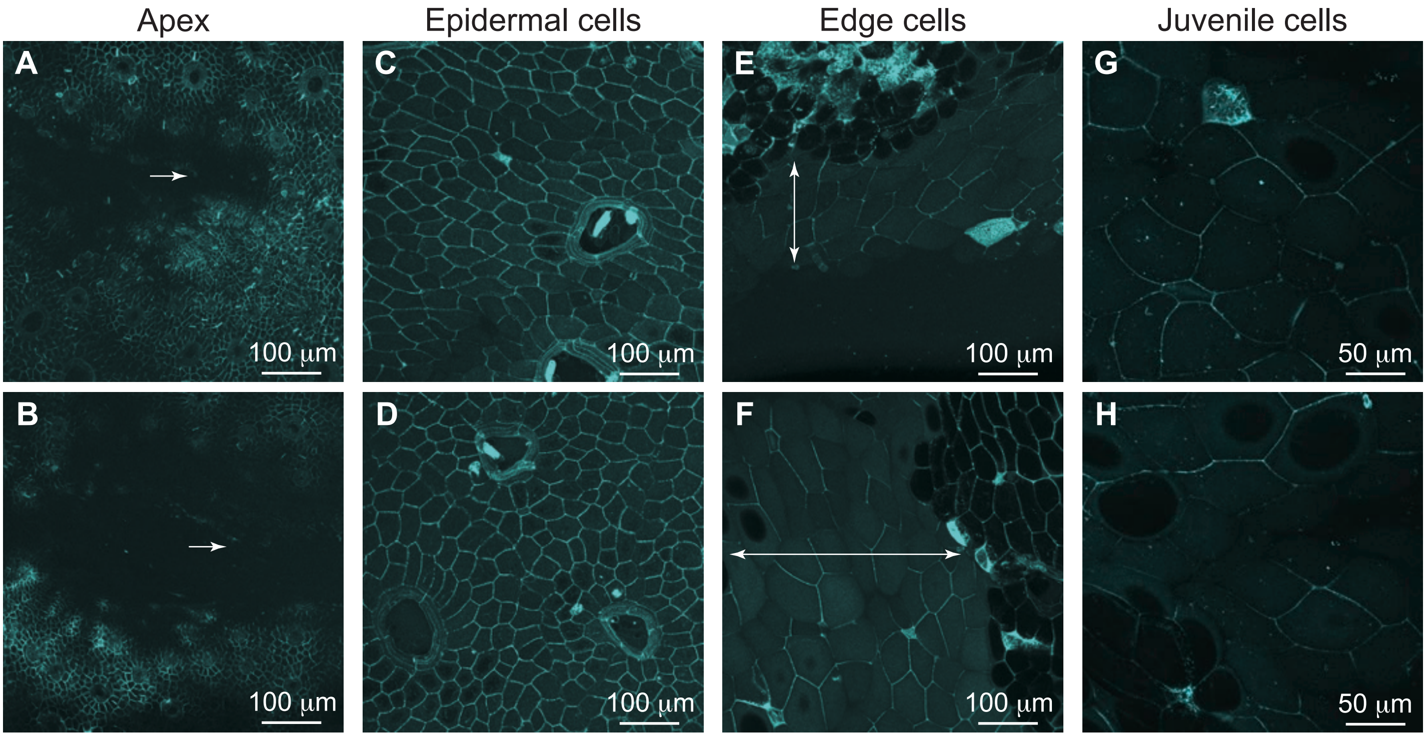


**Fig S9. Aniline blue staining of *M. polymorpha* thalli.**

Maximum projection confocal images of 14-day-old thalli stained with aniline blue. (A, B) Apical notch region. White arrows mark the position of the apical cell, with strong signals observed around the apical notches. (C, D) Epidermal cells. Strong signals are detected in most cells. (E, F) Edge cell areas, marked by double-headed arrows. (G, H) Juvenile cells located at the tail of the thalli. Scale bars are provided in the figures.
